# Targeting Distinct Cell Cycle Nodes Overcomes KRAS/RAS Inhibitor Resistance

**DOI:** 10.64898/2026.03.10.710937

**Authors:** Vishnu Kumarasamy, Jianxin Wang, Edwin Yau, Ethan V. Abel, Agnieszka K. Witkiewicz, Erik S. Knudsen

**Author notes:** **<u>Correspondence</u>:** Erik S. Knudsen, Department of Molecular and Cellular Biology, Roswell Park Comprehensive Cancer Center, Elm and Carlton Streets, Buffalo, NY 14263, Vishnu Kumarasamy, Department of Molecular and Cellular Biology, Roswell Park Comprehensive Cancer Center, Elm and Carlton Streets, Buffalo, NY 14263.

## Abstract

Activating mutations in KRAS drive pancreatic ductal adenocarcinoma (PDAC) and non-small cell lung cancer (NSCLC). Although mutant-selective KRAS inhibitors and pan-RAS inhibitors provide clinical benefits, the development of resistance limits durable response. Transcriptomic and proteomic analyses reveal that, despite effective suppression of mutant KRAS signaling, resistant cells sustain cell cycle progression. Distinct orthogonal mitogenic pathways are engaged in a context-dependent manner to bypass KRAS inhibition. While these pathways can be broadly inhibited using the pan-RAS-ON inhibitor RMC-6236, cells remained capable of developing acquired resistance where cell proliferation is uncoupled from RAS signaling. Combinatorial drug screens and genome-wide CRISPR-Cas9 screens reveal that perturbing cell cycle nodes via targeting cyclin dependent kinases CDK4/6 and CDK2 could restore sensitivity to KRAS/RAS inhibitors. Co-targeting CDK4/6 induces G1 arrest and suppresses E2F-regulated proteins across all resistant models. In contrast, co-targeting CDK2 exerts a broader effect by impairing DNA replication, inducing G2 arrest, preventing mitotic entry, and yielding a more durable cytostatic response that delays cellular outgrowth after drug withdrawal. Finally, concurrent inhibition of KRAS with either CDK4/6 or CDK2 yields durable tumor control *in vivo* in xenografts derived from acquired resistant models. In conclusion, our findings identify sustained cell cycle activity as a defining feature of resistance to KRAS-directed therapies and establish cell cycle co-targeting as an effective strategy to overcome KRAS/RAS inhibitor resistance.

## Introduction

KRAS belongs to the RAS family of small GTPases that regulate fundamental cellular processes including proliferation, differentiation, and survival. In response to growth factor signaling, KRAS transduces signals from receptor tyrosine kinases to activate downstream effector pathways, particularly the RAF-MEK-ERK and PI3K-AKT-mTOR, to promote controlled cell proliferation. These KRAS driven pathways converge to express cyclin D1, which initiates cell cycle progression via activating CDK4 and CDK6 kinases [1]. Oncogenic mutations in KRAS are among the most prevalent drivers of tumorigenesis across multiple cancer types, occurring in approximately 90% of pancreatic ductal adenocarcinoma (PDAC), 43% of colorectal cancer, and 32% of non–small cell lung cancer (NSCLC) [2, 3]. The most frequent alterations are missense mutations at codon 12, where glycine is substituted with aspartate (G12D), cysteine (G12C), or valine (G12V) [2]. These mutations lock KRAS in a constitutively active GTP-bound state, leading to persistent mitogenic signaling independent of upstream regulation and thereby promoting malignant transformation [4].

The oncogenic role of KRAS has been demonstrated in preclinical mouse models, where genetic ablation of KRAS suppresses cell proliferation and regresses tumor growth in NSCLC and PDAC [5]. These findings positioned KRAS as an effective therapeutic target; however, developing pharmacological inhibitors has remained challenging [6]. Recently mutant-specific KRAS and pan-RAS inhibitors were developed that have shown encouraging clinical activity across RAS-driven tumors [7]. Sotorasib (AMG510) and adagrasib (MRTX849) were the first KRAS^G12C^ inhibitors that were clinically approved for the treatment of NSCLC after demonstrating significant improvements in progression-free survival [8, 9]. Currently, a KRAS^G12D^-selective inhibitor, zoldonrasib (RMC-9805) and a pan-RAS-ON inhibitor, daraxonrasib (RMC-6236) are being evaluated in phase III clinical trials and have shown promising efficacy in multiple RAS-driven cancers including PDAC and NSCLC [10, 11]. However, despite these advances, the emergence of acquired resistance remains a major concern, as it limits the durability of the therapeutic response. Preclinical studies have shown that tumors rapidly relapse following drug withdrawal, and in clinical settings, many patients experience disease progression after initial response to KRAS inhibitors [12–14]. Accumulating evidence indicates that resistance frequently arises through compensatory activation of other RAS-isoforms and engagement of parallel mitogenic signaling pathways that restore proliferative capacity [15, 16].

Co-targeting compensatory parallel pathways have emerged as a promising strategy to overcome resistance to KRAS inhibitors [17]. For example, combined inhibition of EGFR with Sotorasib (AMG510) has demonstrated clinical benefit in colorectal cancer; however, the efficacy of this approach in other tumor types is not clear [18, 19]. A key challenge of this strategy is precisely identifying the compensatory pathway to be co-targeted, as these signaling adaptations are highly heterogeneous across tumors. In some contexts, targeting a single parallel pathway may be insufficient to restore therapeutic sensitivity. These challenges underscore the need to better understand the downstream effectors of KRAS signaling that are differentially impacted in resistant models and may represent potential therapeutic targets. In this study, we developed models of acquired resistance to KRAS inhibition and systematically investigated the biological outcomes that sustain proliferation in resistant cells, with the goal of identifying more broadly effective therapeutic vulnerabilities.

## Methods

AsPC-1 and H358 cells were purchased from the American Type Culture Collection (ATCC) and cultured in RPMI-1640 medium supplemented with 10% fetal bovine serum (FBS) and 1% antibiotic–antimycotic (Thermo Fisher Scientific). UM53 is a patient derived PDAC cell line, obtained from Dr. Ethan Abel’s laboratory (Roswell Park Comprehensive Cancer Center, Buffalo, NY) and maintained in RPMI-1640 medium supplemented with 10% FBS and 1% antibiotic–antimycotic [20]. MiaPaCa-2 cells were purchased from the ATCC and cultured in Dulbecco’s Modified Eagle Medium (DMEM) supplemented with 10% FBS and 1% antibiotic–antimycotic. The patient-derived RS4774 cell line was cultured in DMEM supplemented with 20 ng/mL epidermal growth factor (EGF), 10% FBS, and 1% antibiotic–antimycotic. Palbociclib, MRTX849, and MRTX1133 were purchased from ChemieTek (Indianapolis, IN). AMG-510 (sotorasib), RMC-6236, and RMC-9805 were purchased from MedChemExpress (Monmouth Junction, NJ). Compounds were reconstituted in dimethyl sulfoxide (DMSO) to generate stock solutions. INX-315 was provided by Incyclix Bio. The ARTNet drug library comprising a customized panel of therapeutic agents was purchased from Selleck Chemicals (Houston, TX). All cell lines were confirmed to be mycoplasma-free based on DAPI staining and PCR. ASPC-1, H358 and MiaPaCa-2 cells were authenticated by short tandem repeat (STR) profiling.

### Generation of acquired-resistance cell lines

MiaPaCa-2 and H358 cells were initially exposed to a low concentration of MRTX849 (10 nM) and cultured until confluence. The concentration of MRTX849 was increased two-fold every two passages, and cells were continuously maintained in drug-containing medium until the concentration reached 1 µM. Similarly, AsPC-1 cells were cultured in gradually escalating concentrations of MRTX1133 following the same protocol. The resulting resistant cell lines were designated MiaPaCa-2-MR, H358-MR, and AsPC-1-MR. MiaPaCa-2-MR cells were subsequently maintained in a low maintenance dose of MRTX849 (50 nM). However, during experimental procedures, the cells were grown in drug-free medium. To generate MiaPaCa-2-RR cells, parental MiaPaCa-2 cells were cultured in escalating concentrations of RMC-6236 until a final concentration of 200 nM was achieved. Resistance was validated by comparing the effects of KRAS inhibitors on cell proliferation in resistant versus parental cell lines.

### Generation of isogeneic stable cell lines

CRISPR–Cas9–mediated deletion of CDK2 was performed using the pL-CRISPR-EFS-tRFP vector encoding the guide RNA sequence 5′-GCATGGGTGTAAGTACGAACA-3′ targeting CDK2. RFP-positive cells were isolated by fluorescence-activated cell sorting (FACS) using a FACSAria II cell sorter (BD Biosciences), as previously described [21]. Single-cell clones were obtained by subclonal selection, and CDK2 deletion was confirmed by western blotting.

### Cell proliferation assay

Live-cell imaging systems, including IncuCyte S3 (Sartorius), CellCyte X (Cytena), and Cytation (BioTek), were used to monitor cell proliferation in real time. Cell lines were engineered to stably express H2B-GFP and seeded in 96-well tissue culture plates at a density of 1,000–2,500 cells per well, depending on the cell type. These systems quantified the number of H2B-GFP-positive nuclei over time to measure cell proliferation. Fold change in cell number was calculated by normalizing to the initial time point. This approach has been reported in our previous studies [21–23]

### siRNA-mediated knockdown

Cell lines were reverse transfected with gene-specific RNAi that targeted *CDK2*, as described previously [21, 24]. On-target plus human RNAi for *CDK2* was purchased from Horizon Discovery (Cat# L-003236-00-0005). Following transfection, cells were treated with MRTX849 and RMC-6236, and cell proliferation was monitored using live-cell imaging.

### Transcriptome analysis

MiaPaCa-2-WT and MiaPaCa-2-MR cells were treated with MRTX849 at a concentration of 100 nM up to 48 hours. Total RNA was extracted using the Qiagen RNeasyplus kit and the RNA quality was evaluated using the RNA6000 Nano assay and with the Agilent 2200 TapeStation (Agilent, CA, USA) [22]. Th entire protocol was described in our previous studies [21, 22]. Complementary DNA (cDNA) was synthesized from purified RNA using random hexamer priming to capture non-ribosomal transcripts. Sequencing libraries were prepared using the DriverMap Human Genome-Wide Gene Expression Profiling Kit (hDM18Kv3; Cellecta Inc.) following the manufacturer’s protocol. Anchor PCR amplification was performed under the following conditions: initial denaturation at 95 °C for 5 min; 15 cycles of 95 °C for 30 s, 68 °C for 1 min, and 72 °C for 1 min; followed by a final extension at 72 °C for 10 min. PCR products were purified using AMPure XP SPRI beads (Beckman Coulter) at a 1:1 sample-to-bead ratio, and library concentrations were quantified using the Qubit dsDNA High Sensitivity Assay Kit (Thermo Fisher Scientific) [21, 22]. Target-enriched RNA-seq libraries were sequenced on an Illumina NextSeq 500 platform using the NextSeq 500/550 High Output v2 kit (75-cycle configuration) according to the manufacturer’s instructions. Sequencing reads were aligned to the human reference genome using STAR aligner (v2.7.10b), and gene-level read counts were generated. Differential gene expression analysis was performed using edgeR, and significantly regulated genes were identified based on statistical significance and log_₂_ fold change thresholds.

### CRISPR screening

The Toronto Knockout (TKO) CRISPR library version 3 was packaged into lentiviral particles and were used to infect the UM53 cell lines as described in previous studies [21, 22, 25]. The positive clones were further expanded in the absence and presence of MRTX849 (500 nM) and RMC-6236 (7.5 nM). Following the fifth passage, the cells were harvested and subjected to genomic DNA extraction using the Wizard Genomic DNA purification kit (Promega, A1120). For downstream analysis, genomic DNA was isolated and subjected to a two-step PCR amplification to enrich sgRNA sequences integrated into the genome and to incorporate Illumina TruSeq adapters with i5 and i7 sample indices, as previously described [22]. The resulting PCR products were purified and used to generate sequencing libraries.

Libraries were sequenced on an Illumina NextSeq platform to obtain single-end reads in FASTQ format. Adapter sequences and low-quality bases were trimmed using Trim Galore (v0.6.7; https://github.com/FelixKrueger/TrimGalore) [22]. Trimmed reads were then processed using the MAGeCK pipeline to quantify sgRNA read counts for each sample. The resulting count matrix was used for downstream analysis, including identification of genes associated with drug response using the DrugZ algorithm.

### Proteomics analysis

To determine differential protein expression between the untreated and MRTX849-treated cells, whole cell extracts were prepared using the IP-lysis buffer 20 mM Tris-HCl, pH 8.0, 2mM EDTA, 137 mM NaCl, 1% NP-40) in the presence of Halt Protease inhibitor cocktail and 1 mM PMSF [21]. Extracted proteins were digested prior to LC–MS analysis. Briefly, protein lysates were reduced with 200 mM dithiothreitol (DTT) at 58 °C for 30 min, followed by alkylation with 500 mM iodoacetamide at 37 °C for 30 min. Proteins were then precipitated using pre-chilled acetone and subjected to trypsin digestion at 37 °C for 6 h. Digestion was terminated by acidification with 1% formic acid. Peptides were analyzed by LC–MS, and differential protein expression was determined based on log_₂_ fold change in peptide abundance between treatment conditions.

### Western blotting

Cells were lysed, and whole-cell extracts were prepared using RIPA lysis buffer (10 mM Tris-HCl, pH 8.0, 1 mM EDTA, 150 mM NaCl, 1% Triton X-100, 0.1% sodium deoxycholate, 0.1% SDS) supplemented with Halt protease inhibitor cocktail and 1 mM PMSF [21, 22]. Primary antibodies against phospho-ERK (Thr202/Tyr204; Cat# 9101), ERK (Cat# 9102), phospho-RB (Ser807/811; D20B12; Cat# 8516S), Cyclin B1 (D5C10; Cat# 12231S), CDK2 (78B2; Cat# 2546S), and CDK1 (Cat# 77055) were purchased from Cell Signaling Technology (Danvers, MA). Antibodies against Cyclin A (Cat# AF5999) and β-actin (Cat# MAB8929) were purchased from R&D Systems (Minneapolis, MN). Primary antibodies were diluted 1:1000 in antibody dilution buffer (5% BSA in 25 mM Tris-HCl, pH 7.5, 150 mM NaCl, 0.1% Tween-20). HRP-conjugated anti-mouse, anti-rabbit, and anti-goat secondary antibodies were purchased from Santa Cruz Biotechnology (Dallas, TX). Protein bands were detected using SuperSignal West Femto chemiluminescent substrate (Thermo Fisher Scientific) and imaged using a Bio-Rad imaging system.

#### In vitro kinase reactions

The catalytic activity of CDK2 was assessed using *in vitro* kinase reactions [21, 23, 26]. Cells were lysed to extract proteins using the kinase assay lysis buffer (50 mM HEPES-KOH pH 7.5, 1mM EDTA, 150 mM NaCl, 1mM DTT, 100% Glycerol and 0.1% Tween-20). CDK2 was immunoprecipitated using anti-CDK2 antibody (SC-6248). The kinase reaction was carried out in the kinase assay buffer (40 mM Tris-HCl pH 8.0, 20 mM MgCl_2_, 0.1 mg/ml BSA and 50 µM BSA) in the presence of 500 µM ATP. The RB C-terminal peptide was used as an exogenous substrate [27].

### Cell cycle analysis

To assess the distribution of cells across different cell cycle phases, we followed the protocol that was published in our previous study [21]. Cells were fixed in ice-cold 70% ethanol overnight at −20 °C. Fixed cells were washed with 1× PBS and incubated with RNase A (200 µg/mL) and propidium iodide (PI; 40 µg/mL) prior to analysis. For BrdU incorporation analysis, cells were pulsed with BrdU for 3 h and fixed in ice-cold 70% ethanol overnight at −20 °C. Fixed cells were denatured in denaturation buffer (2 N HCl, 0.5% Triton X-100) for 30 min at room temperature, followed by neutralization with 0.1 M sodium tetraborate (pH 8.5). Cells were then washed with IFA buffer (1% BSA in 1× PBS) and incubated with FITC-conjugated anti-BrdU antibody for 1.5 h at room temperature. After washing with IFA buffer, cells were resuspended in PBS containing RNase A and PI. Cell cycle distribution and BrdU incorporation were analyzed using a BD LSRFortessa flow cytometer (BD Biosciences), and data were analyzed using FCS Express software.

### Mice and patient derived xenografts

NSG mice were bred and maintained at the Roswell Park Comprehensive Cancer Center animal facility. All animal procedures, including housing, tumor implantation, drug administration, and euthanasia, were approved by the Roswell Park Comprehensive Cancer Center Institutional Animal Care and Use Committee (IACUC) and conducted in accordance with the NIH Guide for the Care and Use of Laboratory Animals. MiaPaCa-2-MR xenografts were developed in 10-week-old NSG male mice by subcutaneously injecting 5×10^6^ cells/mouse. Resulting tumors were excised and serially passaged subcutaneously. The mice bearing MiaPaCa-2-MR xenografts were randomized into 6 groups: Vehicle (n=9) and palbociclib (n=4), MRTX849 (n=5) and palbociclib+MRTX849 (n=4), INX-315 (n=5), INX-315+MRTX849 (n=6). Patient derived lung cancer xenografts, RS4774 PDX were implanted subcutaneously into 10-week-old male NSG mice and were randomized into 4 groups: Vehicle (n=7) and palbociclib (n=4), MRTX849 (n=4) and palbociclib+MRTX849 (n=5). Palbociclib was formulated in lactate buffer (pH 4.0) and administered by oral gavage at 100 mg/kg. MRTX849 was formulated in 5% DMSO, 40% PEG300, and 55% saline and administered by oral gavage at 15 mg/kg. INX-315 was formulated in 100% polyethylene glycol-400 (PEG400) and administered daily by oral gavage at 100 mg/kg, as previously described [21]. Tumor growth was monitored every other day using digital calipers, and tumor volume was calculated using standard methods. The maximum tumor volume permitted under our institutional guidelines was 2,000 mm³, at which point mice were euthanized. Mice that died during the course of treatment were excluded from the analysis

### Histological analysis

Tumor tissues that were excised from the mice were fixed in 10% Formalin followed by processing and paraffin embedding as described in our previous study [21]. The embedded tissues were serially sectioned at 4-6 µm using the standard procedures and were subjected to Hematoxylin and Eosin (H&E) staining [22]. RB phosphorylation was determined by immunohistochemical staining using the pRB (S807/811) antibody (Cat # 8516) (Cell signaling Technologies). Slides were scanned using the Vectra Polaris Instrument as described before [21, 22].

## Results

### Development of acquired resistance to mutant-specific KRAS inhibitors

Previous studies have demonstrated that KRAS^G12C^-driven cell lines H358 (NSCLC) and MiaPaCa-2 (PDAC) and the KRAS^G12D^-driven cell line AsPC-1 (PDAC) are highly sensitive to mutant-specific KRAS inhibitors such as AMG510, MRTX849 and MRTX1133 [22, 28, 29]. Acquired resistance was modeled *in vitro* by subjecting the H358, MiaPaCa-2 and ASPC-1 cells to prolonged exposure to the KRAS^G12C^ inhibitor MRTX849 and the KRAS^G12D^ inhibitor MRTX1133, with gradual dose escalation over a period of up to 6 months. Resistant derivatives that emerged under continuous drug pressure were referred to as H358-MR, MiaPaCa-2-MR and AsPC-1-MR cells. To validate the resistance phenotype, the anti-proliferative effect of the KRAS inhibitors on the parental and acquired resistant cell lines were evaluated using live cell imaging. The parental cell lines, MiaPaCa-2, ASPC-1 and H358 displayed a robust suppression of cell proliferation in a dose-dependent manner (Fig. 1A). In contrast, the resistant derivatives, MiaPaCa-2-MR, AsPC-1-MR and H358-MR cells continued to proliferate despite the presence of KRAS inhibitors (Fig. 1B). A patient-derived lung cancer cell line, RS4774, was also developed from a in individual treated at Roswell Park that developed disease progression post treatment with a KRAS^G12C^ inhibitor, AMG510 (NCT03600883) [30]. The established cell line harbored a single allele mutation in the *KRAS* gene (G12C) and retained the resistant phenotype to MRTX849 as indicated by the modest effect on proliferation (Fig. 1C, S1A). Additionally, another patient-derived PDAC cell line UM53 that harbors the KRAS^G12C^ mutation displayed intrinsic resistance to the KRAS inhibitor MRTX849, which is consistent with our previous study (Fig. 1C) [22, 31].

**Figure 1:**
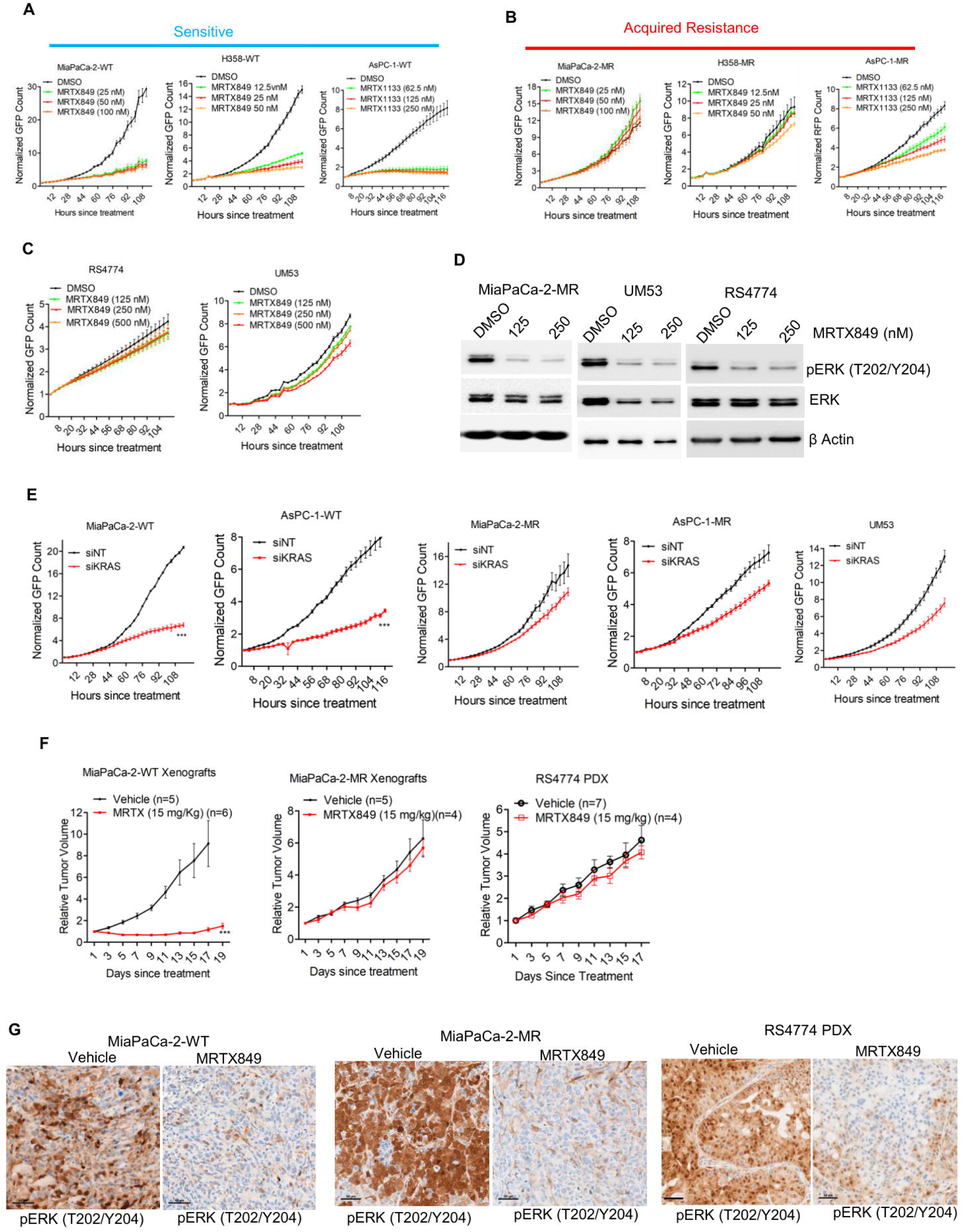
Single agent efficacy of mutant specific KRAS inhibitors: (A) Dose-dependent effect of MRTX849 on the proliferation of MiaPaCa-2 and H358 cells based on live cell imaging. AsPC-1 cells were treated with different concentrations of MRTX1133, and the cell growth was monitored. Error bars represent mean and SD from triplicates. Experiments were done at three independent times. (B) Effect of the mutant-specific KRAS inhibitors, MRTX849 and MRTX1133 on the proliferation of MiaPaCa-2-MR, H358-MR and AsPC-1-MR cells that have developed acquired resistance following long-term selection. Mean and SD were determined from triplicates, and the experiments were carried out 3 independent times. (C) Live cell imaging to examine the effect of MRTX849 on the proliferation of UM53 and RS4774 cell lines. Error bars represent mean and SD from triplicates. Experiments were done at three independent times. (D) Western blot analysis on MiaPaCa-2-MR, UM53 and RS4774 cells to evaluate the effect of MRTX849 on ERK phosphorylation following 48-hour exposure at the indicated concentration. (E) Differential effects of KRAS knockdown on the proliferation of MiaPaCa-2-WT, AsPC-1-WT, MiaPaCa-2-MR, AsPC-1-MR and UM53 cells. Error bars indicate mean and SD from triplicates. *** represents p<0.0001 as determined by 2-way ANOVA. Experiments were done at three independent times. (F) *In vivo* efficacy of MRTX849 on xenografts derived from MiaPaCa-2-WT and MiaPaCa-2-MR cells and on the tumor growth of RS4774 PDX. Error bars represent mean and SEM. *** represents p<0.0001 as determined by 2-way ANOVA. (G) Representative images of immunohistochemical staining on the vehicle- and MRTX849-treated tissues to determine the phosphorylation status of ERK. Scale bar represents 50 microns.

To determine whether resistance was mediated by impaired drug effect on mutant KRAS, we examined the effect of KRAS inhibition on ERK phosphorylation, a key downstream effector of RAS signaling [32]. In all resistant models, treatment with KRAS inhibitors resulted in suppression of ERK phosphorylation, indicating effective pharmacologic inhibition of mutant KRAS activity (Fig. 1D). Despite sustained pathway inhibition, resistant cells continued to proliferate, suggesting that their growth is no longer dependent on mutant KRAS signaling. Consistent with this observation, genetic depletion of KRAS using gene-specific siRNA significantly inhibited proliferation in parental MiaPaCa-2 and AsPC-1 cells (Fig. 1E). In contrast, KRAS depletion had minimal impact on proliferation in the corresponding acquired resistant models (MiaPaCa-2-MR and AsPC-1-MR), as well as in the intrinsically resistant UM53 cells (Fig. 1E). To further examine that the resistant phenotype is not drug-specific but instead reflects resistance to KRAS pathway inhibition, we evaluated the effects of additional mutant-selective inhibitors, AMG510 and RMC-9805 that inhibit KRAS^G12C^ and KRAS^G12D^ respectively. AMG510 more potently suppressed proliferation in MiaPaCa-2-WT cells compared with MiaPaCa-2-MR cells (Fig. S1B). Likewise, RMC-9805 showed greater antiproliferative activity in AsPC-1-WT cells than in AsPC-1-MR cells (Fig. S1C), demonstrating that resistance extends across distinct mutant-selective KRAS inhibitors. Together, these findings demonstrate that resistance is mediated through functional decoupling of cell proliferation from mutant KRAS activity.

Since our previous studies indicated that resistance to KRAS inhibitors can be attenuated *in vivo*, we evaluated whether the acquired resistance phenotype observed in cell culture could be maintained *in vivo* [22]. To this end, we compared the efficacy of KRAS inhibition in xenografts derived from parental MiaPaCa-2 and acquired resistant MiaPaCa-2-MR cells. Consistent with our *in vitro* findings, treatment with MRTX849 resulted in significant tumor growth inhibition in MiaPaCa-2-WT xenografts, whereas MiaPaCa-2-MR xenografts exhibited only a modest response at the indicated dose (Fig. 1F). Similarly, the RS4774 PDX retained resistance to MRTX849 as demonstrated by continuous tumor growth despite KRAS inhibition (Fig. 1F). To confirm effective target engagement *in vivo*, ERK phosphorylation was assessed by immunohistochemistry in tumor tissues. MRTX849 treatment resulted in the suppression of ERK phosphorylation in the sensitive MiaPaCa-2-WT and resistant MiaPaCa-2-MR and RS4774 xenografts, indicating effective inhibition of KRAS signaling despite differential tumor growth responses (Fig. 1G). In conclusion, acquired resistance limits the anti-proliferative efficacy of KRAS inhibitors, both *in vitro* and *in vivo*.

### Cellular mechanisms underlying acquired resistance to KRAS inhibition

To investigate the molecular outcomes following KRAS inhibition that defines differential response between the sensitive and resistant models, we performed global transcriptomic and proteomic analysis in MiaPaCa-2-WT and MiaPaCa-2-MR cells following treatment with MRTX849. Gene set enrichment analysis (GSEA) of differentially expressed genes and proteins revealed that the top pathways that were significantly impacted were the E2F-targets and G2/M checkpoint, which are involved in the cell cycle machinery; these pathways were more significantly downregulated in the parental MiaPaCa-2-WT cells compared with the acquired resistant MiaPaCa-2-MR cells (Fig. 2A, S2A). The magnitude of repression of individual cell cycle genes and proteins was more pronounced in the MiaPaCa-2-WT than in the MiaPaCa-2-MR cells (Fig. 2B). Given that E2F-driven transcription is regulated via RB phosphorylation, we carried out biochemical analysis to evaluate its phosphorylation status. KRAS inhibition led to robust RB dephosphorylation in sensitive models (H358-WT, MiaPaCa-2-WT, ASPC-1-WT) whereas both acquired (H358-MR, MiaPaCa-2-MR, ASPC-1-MR) and intrinsically resistant models (UM53 and RS4774) retained RB phosphorylation (Fig. 2C). Expression of the E2F-regulated cell cycle protein cyclin A correlated with RB phosphorylation status across models (Fig. 2C). Overall, our data indicates that sustained cell cycle activity represents a key cellular outcome associated with resistance to KRAS inhibition.

**Figure 2:**
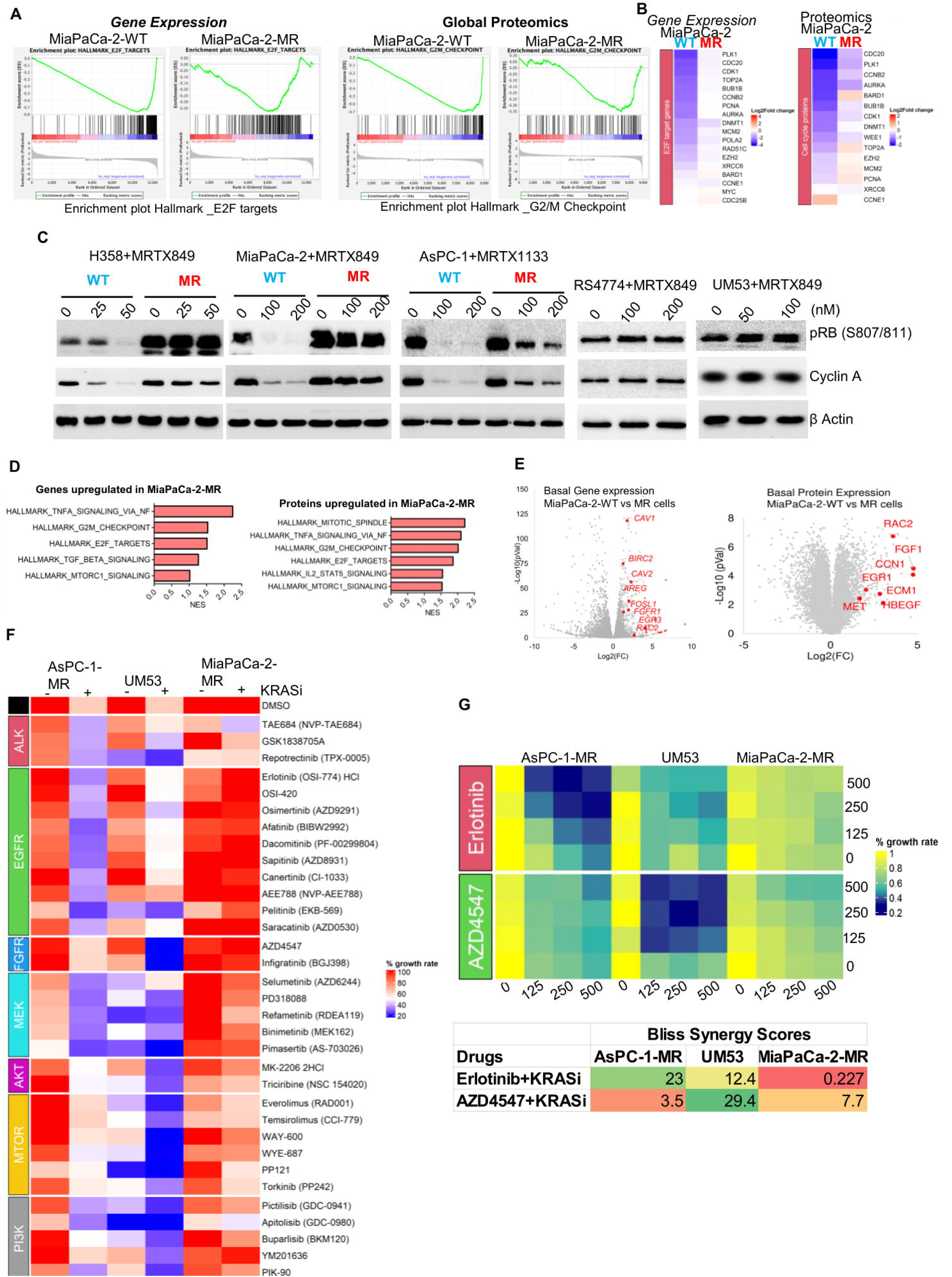
Cellular outcomes that define differential responses to KRAS inhibitors: (A) GSEA analysis illustrating the differential effect of MRTX849 on the genes associated with E2F target pathway and the proteins associated with G2/M checkpoint in cell cycle machinery. (B) Heat map represents the differential effect of MRTX849 in downregulating the indicated genes and proteins that are involved in cell cycle from MiaPaCa-2-WT and MiaPaca-2-MR cells following 48-hour treatment. (C) Biochemical analysis on the indicated sensitive (WT) and resistance cell lines to compare the effect of MRTX849 on RB phosphorylation and cyclin A expression at different doses after treating the cells for 48 hours. (D) GSEA analysis highlighting pathways associated with genes and proteins that are differentially expressed between MiaPaCa-2-MR and MiaPaCa-2-WT cells. (E) Volcano plots of differentially expressed genes and proteins in MiaPaCa-2-MR versus MiaPaCa-2-WT cells. (F) Heat map illustrating the effects of indicated targeted therapeutic agents on normalized fold change in cell growth in combination with DMSO or mutant-selective KRAS inhibitors. (G) Synergistic interactions between KRAS inhibitors and erlotinib or AZD4547 in AsPC-1-MR, UM53, and MiaPaCa-2-MR cells. Heat maps depict normalized fold change in growth rate following treatment with increasing concentrations of KRAS inhibitors in combination with erlotinib or AZD4547. Bliss synergy scores were calculated using the SynergyFinder online platform.

To identify molecular pathways that may contribute to resistance to KRAS inhibition, we compared basal gene and protein expression profiles between parental MiaPaCa-2-WT and acquired resistant MiaPaCa-2-MR cells. Gene set enrichment analysis (GSEA) of genes and proteins differentially expressed in MiaPaCa-2-MR cells relative to MiaPaCa-2-WT cells revealed significant enrichment of multiple growth-associated signaling pathways, including TNFα signaling, mTOR signaling, and mitotic cell-cycle programs (Fig. 2D). At the individual gene and protein level, MiaPaCa-2-MR cells exhibited increased expression of several growth factors and receptor-associated components, including *CAV1, AREG, FGF1,* and *HBEGF* (Fig. 2E). The elevated expression of these ligands and associated proteins suggests the potential involvement of receptor tyrosine kinase (RTK)-mediated signaling pathways that may contribute to drive resistance. In addition, downstream RAS pathway effectors, including RAC2 and FOSL1, were upregulated at both the transcript and protein levels in MiaPaCa-2-MR cells (Fig. 2E). Together, these findings indicate that acquired resistance to KRAS inhibition is associated with engagement of parallel growth-promoting signaling networks that bypass KRAS dependency and sustain cell proliferation.

As a complementary approach to demonstrate the involvement of parallel mitogenic signaling pathways contributing to KRAS inhibitor resistance, we performed a combinatorial drug screen to determine whether co-targeting different pathways could restore sensitivity in resistant cells. The resistant AsPC-1-MR, MiaPaCa-2-MR, and UM53 cells were pretreated with mutant-specific KRAS inhibitor, subsequently exposed to a targeted drug library and combination effects were assessed based on cell growth inhibition. This analysis revealed that co-targeting distinct signaling pathways enhanced the efficacy of KRAS inhibition in a context-dependent manner. As depicted in the heatmap, co-targeting EGFR selectively enhanced the anti-proliferative effect of KRAS inhibition in AsPC-1-MR cells, while producing only modest effects in UM53 cells (Fig. 2F). In contrast, inhibition of FGFR or mTOR pathways was more effective in UM53 cells, with limited impact in AsPC-1-MR cells (Fig. 2F). Notably, MiaPaCa-2-MR cells were broadly refractory to most agents targeting mitogenic signaling pathways, suggesting that inhibition of a single parallel pathway is insufficient to overcome resistance in this context (Fig. 2F). Synergy analyses validated these screening results, demonstrating selective synergistic interactions between EGFR and KRAS inhibition in AsPC-1-MR cells, and between FGFR and KRAS inhibition in UM53 cells (Fig. 2G, Fig. S3A). Additionally, MiaPaCa-2-MR cells showed only modest responses to either combination strategy (Fig. 2G, S3A). Collectively, these data indicate that while co-targeting parallel signaling pathways can restore sensitivity to KRAS inhibitors, the magnitude and durability of response are highly heterogeneous and strongly dependent on cellular context.

### Impact of pan-RAS-ON inhibition in KRAS inhibitor resistant models

Given the activation of multiple parallel mitogenic pathways in resistant models, we next investigated a therapeutic strategy aimed at broadly suppressing these pathways. To test this, we evaluated the pan-RAS-ON inhibitor RMC-6236, a recently developed active-RAS inhibitor with emerging clinical activity in RAS-driven tumors [11]. Across the resistant models, RMC-6236 yielded a robust cytostatic effect, which was significantly more potent than the mutant specific inhibitors (Fig. 3A). Consistent with this observation, the EC_₅₀_ values for RMC-6236 were significantly lower than those for MRTX849 and MRTX1133 across resistant models (Fig. 3B). To investigate the cellular processes differentially affected by mutant-specific KRAS versus pan-RAS inhibition, we performed gene expression analysis in AsPC-1-MR cells. Gene set enrichment analysis revealed that RMC-6236 treatment led to a stronger suppression of mitogenic pathways such as TNFα and MTORC signaling and the cell cycle machinery as compared to the effect of MRTX1133 (Fig. 3C). The magnitude of repression of the genes involved in the MEK signaling, MTOR signaling and cell cycle progression are more prominent in the presence of RMC6236 as compared to that with MRTX1133, indicating the enhanced efficacy of RAS-ON inhibition (Fig. 3D). Single-agent efficacy of MRTX1133 in AsPC-1-WT cells was examined as a control, and it suppressed the mitogenic transcriptional program more potently than in AsPC-1-MR cells (Fig. S4A). Biochemical analysis further validated the gene expression data, that in all the KRAS mutant-selective inhibitor-resistant models (AsPC-1-MR, MiaPaCa-2-MR and UM53), RMC-6236 inhibited ERK phosphorylation more effectively than the KRAS inhibitors MRTX1133 and MRTX849 (Fig. 3E). Moreover, RMC-6236 was more effective in yielding dephosphorylation of RB and suppressing the expression of cell cycle proteins such as cyclin A and cyclin B1 as compared to KRAS inhibitors (Fig. 3E). However, long-term colony formation assays in the presence of RMC-6236 revealed heterogeneity in response. Although AsPC-1-MR cells remained highly sensitive, a subpopulation of MiaPaCa-2-MR cells resumed proliferation (Fig. 3F).

**Figure 3:**
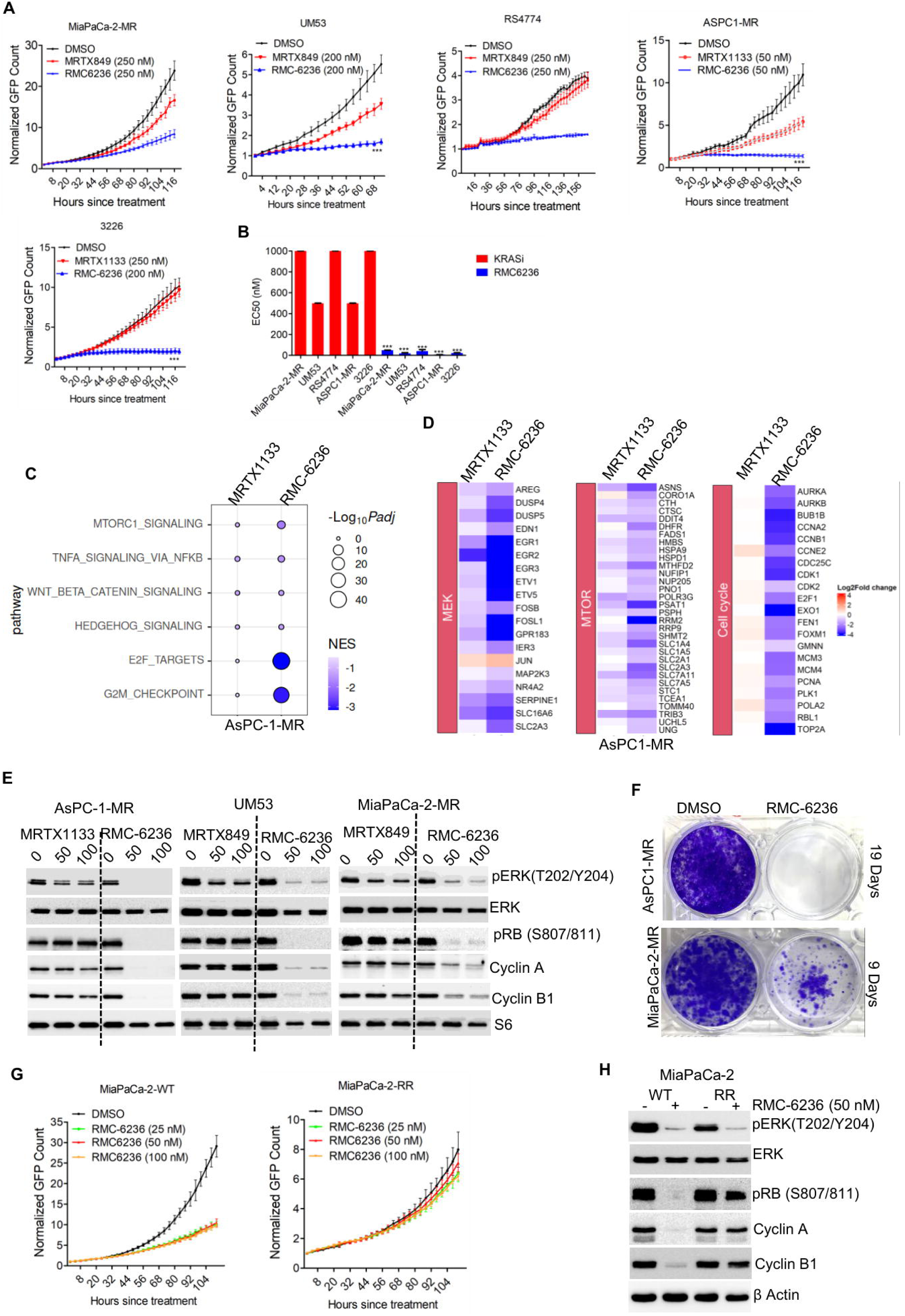
Single agent efficacy of PAN-RAS-ON inhibitor, RMC-6236: (A) Comparison of the antiproliferative effects of mutant-selective KRAS inhibitors and RMC-6236 in the indicated cell lines. Error bars represent mean and SD from triplicates. Experiments were done 3 independent times. *** represents p<0.0001 as determined by 2-way ANOVA. (B) EC_50_ values of mutant specific KRAS inhibitors and RMC-6236 in multiple cell lines. Error bar represents mean and SD from 2 independent experiments. *** represents p<0.0001 as determined by unpaired student t-test. (C) GSEA analysis highlighting the top pathways that were differentially impacted in AsPC-1-MR cells following the treatment with MRTX1133 (100 nM) and RMC-6236 (50 nM) for 48 hours. (D) Heatmap depicts the differential expression of the indicated genes associated with MEK, MTOR and cell cycle pathways following the treatment with MRTX1133 and RMC-6236 in AsPC-1-MR cells. (E) Immunoblot analysis comparing the effect of mutant-specific KRAS inhibitors and RMC-62366 on the indicated proteins after exposing the cells to different drug concentrations up to 48 hours. (F) Long term colony formation assay in AsPC-MR and MiaPaCa-2-MR cells following the treatment with RMC-6236. AsPC-1-MR cells were treated with 50 nM RMC-6236 for 19 days and MiaPaCa-2-MR cells were treated 100 nM of RMC-6236 for 9 days. (G) Differential effect of RMC-6236 on the proliferation of MiaPaCa-2-WT and MiaPaCa-2-RR cells following the treatment with different drug concentrations. Error bars represent mean and SD from triplicates. Experiment was done at 3 independent times. (H) Biochemical analysis to compare the effect of RMC-6236 on the indicated proteins between the MiaPaCa-2-WT and MiaPaCa-2-RR cells following 48-hour exposure.

Prolonged exposure of MiaPaCa-2-WT cells to RMC-6236 with gradual dose escalation led to the emergence of an acquired resistant population (MiaPaCa-2-RR). Live-cell imaging confirmed that while RMC-6236 suppressed the proliferation of MiaPaCa-2-WT cells, the impact on MiaPaCa-2-RR cells was quite modest (Fig. 3G). Biochemical analysis demonstrated that ERK phosphorylation was suppressed in both the MiaPaCa-2-WT and MiaPaCa-2-RR cells following treatment with RMC-6236, confirming drug-target interaction (Fig. 3H). However, unlike the MiaPaCa-2-WT cells, RMC-6236 possessed a modest inhibitory effect on RB phosphorylation and the expression of cyclin A and cyclin B1 in MiaPaCa-2-RR cells, indicating that the cell cycle was active despite RAS pathway inhibition (Fig. 3H). Collectively, these findings suggest that although pan-RAS inhibition produces a more potent cytostatic response than mutant-selective KRAS inhibitors, adaptive mechanisms can restore cell cycle progression, limiting the durability of response.

### Pharmacologically targeting the cell cycle overcomes KRAS inhibitor resistance

Given the heterogeneous responses observed with co-targeting signaling pathways and the emergence of acquired resistance to the RAS inhibitor RMC-6236, we sought to identify a strategy that could yield a more uniform response across all resistant models. Interrogating the combinatorial drug interaction screens identified CDK4/6, a key regulator of cell cycle entry, as a common target across all three resistant models (Fig. 4A). Multiple CDK4/6 inhibitors, including palbociclib, abemaciclib, and GT138 (lerociclib), consistently enhanced the anti-proliferative effects of KRAS inhibition in MiaPaCa-2-MR, UM53, and AsPC-1-MR cells (Fig. 4A). Subsequent validation experiments confirmed that combining the CDK4/6 inhibitor palbociclib with KRAS and RAS-ON inhibitors cooperatively suppressed cell proliferation across all resistant models (Fig. S5A). Quantitative synergy analyses demonstrated a strong and comparable synergistic interaction between palbociclib and KRAS or RAS inhibitors across the resistant AsPC-1-MR, UM53, and MiaPaCa-2-MR cells (Fig. 4B). Biochemical analyses revealed that palbociclib in combination with KRAS or RAS inhibitors more effectively suppressed E2F-regulated cell cycle proteins, including cyclin B1 and CDK1, which are critical for cell cycle progression and mitotic entry (Fig. 4D, E) [33]. To determine whether these antiproliferative effects extended beyond 2D cell culture assays, the combination was evaluated in spheroids derived from MiaPaCa-2-MR and UM53 cells. Consistent with the 2D culture findings, the combination of palbociclib and MRTX849 resulted in significantly enhanced suppression of spheroid growth compared to single-agent treatment (Fig. 4E). The acquired resistant lung cancer cell line H358-MR also responded to the combination treatment of palbociclib and MRTX849 in both 2D culture and as spheroids (Fig. S5B, C). Moreover, the co-targeting CDK4/6 also significantly enhanced the efficacy of RMC-6236 to overcome acquired resistance in the MiaPaCa-2-RR cell line (Fig. S5D). Overall, these findings display a cooperative interaction between CDK4/6 and KRAS/RAS inhibitors that leads to suppressed proliferation across different resistant models, yielding a more consistent response than strategies targeting upstream mitogenic signaling.

**Figure 4:**
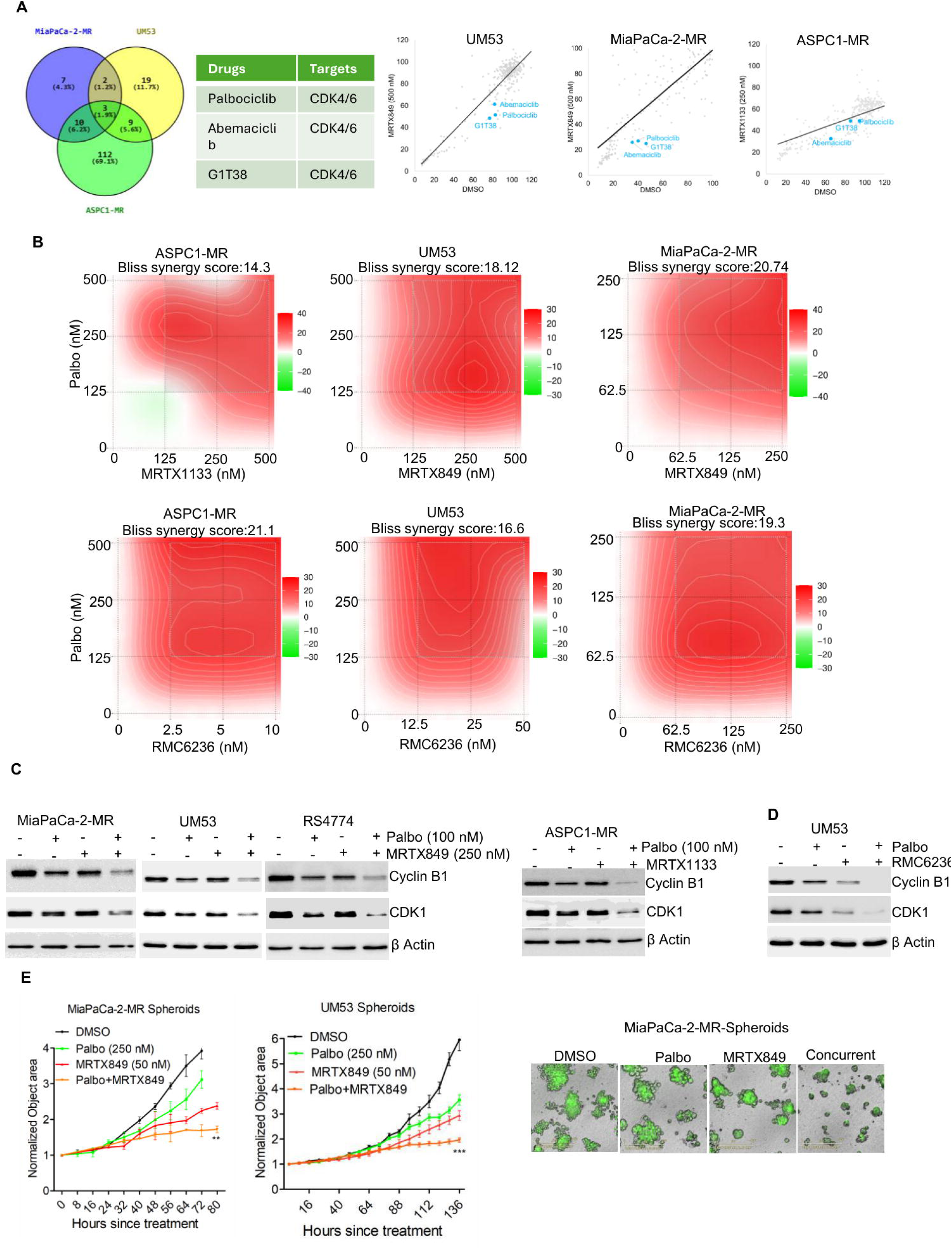
Co-targeting CDK4/6 kinase and KRAS/RAS pathway: Venn diagram showing drugs that enhanced the efficacy of mutant-selective KRAS inhibitors in MiaPaCa-2-MR, AsPC-1-MR, and UM53 cells, identified through combinatorial drug screening. Common hits included the CDK4/6 inhibitors palbociclib, abemaciclib, and G1T38. Correlation plot comparing the fold change in growth rate for individual drugs in combination with DMSO or mutant-selective KRAS inhibitors. (B) Synergistic interactions between palbociclib and KRAS or RAS inhibitors in AsPC-1-MR, UM53, and MiaPaCa-2-MR cells. Isobolograms were generated based on normalized fold change in growth rate following treatment with increasing concentrations of palbociclib in combination with KRAS or RAS inhibitors. Bliss synergy scores were calculated using the SynergyFinder online platform. (C) Western blot analysis in MiaPaCa-2-MR, UM53 and RS4774 following the treatment with palbociclib (100 nM) in combination with MRTX849 (250 nM) up to 48 hours. AsPC-1-MR cells were treated with palbociclib (100 nM) in combination with MRTX1133 (250 nM). (D) Western blotting in UM53 cells to examine the effect of palbociclib (100 nM) in combination with the RAS inhibitor, RMC-6236 (25 nM) on the cell cycle proteins following 48-hour treatment. (E) Effect of Palbociclib in combination with MRTX849 on the growth of spheroids derived from MiaPaCa-2-MR and UM53 cells. Error bars represent mean and SEM from triplicates. *** represents p<0.0001 and ** represents p<0.001 as determined by 2-way ANOVA. Experiments were done at 3 independent times.

### Mechanistic impact of co-targeting RAS pathway and CDK4/6 kinases

Based on prior studies including our own work, it is evident that the mechanistic impact of CDK4/6 inhibitors on cell cycle is imparted through inactivating CDK2, which is a critical kinase involved in allowing the cells to undergo DNA replication and mitosis [23, 34] . Since the combination of palbociclib with KRAS and RAS inhibitors yields a synergistic effect, the impact of this combination on CDK2 kinase activity was assessed. To that end, a live-cell CDK2 sensor was used, which comprises a peptide fragment from CDK2 substrate, DNA helicase B, fused to mCHERRY fluorescent protein as described in previous studies [23, 35]. In active cell proliferation, CDK2 phosphorylates the DHB-mCHERRY substrate, which results in cytoplasmic localization while the inactive kinase localizes the fluorescent signal in the nucleus. As expected, palbociclib in combination with MRTX849 and RMC-6236 significantly decreased the ratio of cytoplasmic to nuclear signal in UM53 cells, indicating an inactive CDK2 when compared to single-agent treatments (Fig. 5A). As a complementary approach, the intracellular CDK2 complex was immunoprecipitated from UM53 and RS4774 cells following treatment with palbociclib in combination with MRTX849 and its catalytic activity was examined *in vitro* using C-terminal peptide fragment from RB as the substrate [27]. The combination treatment resulted in dephosphorylation of RB substrate, further confirming that the CDK2 complex was inactive (Fig. S6A). In the sensitive model MiaPaCa-2, KRAS mutant-selective inhibitor alone was sufficient to inactivate CDK2, inducing a cytostatic response (Fig. S6B). Overall, these data indicate that CDK2 kinase activity functions as a cellular determinant of response to KRAS inhibitors.

**Figure 5:**
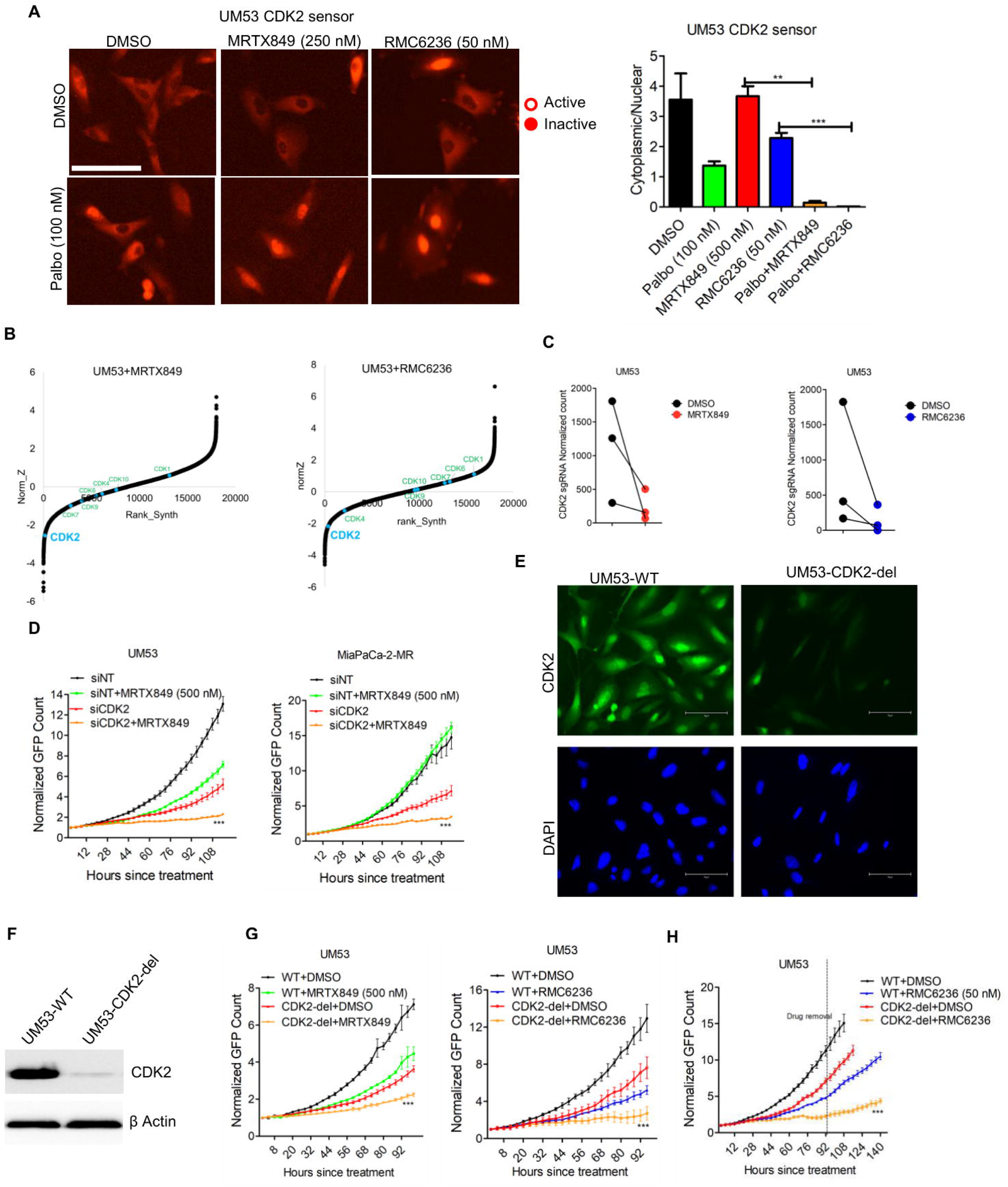
Downstream impact of CDK4/6 in combination with KRAS/RAS inhibitors: (A) Representative images of UM53 cells expressing the CDK2 activity sensor following treatment with palbociclib (100 nM) in combination with MRTX849 (250 nM) or RMC-6236 (25 nM) for up to 72 hours. Bar graphs show the ratio of cytoplasmic to nuclear mCherry signal, corresponding to CDK2 activity, following the indicated treatments. Error bars represent mean and SEM from triplicates. *** represent p<0.0001 and ** represents p<0.001 as determined by unpaired student t-test. Experiment was done at three independent times. (B) DrugZ analysis from UM53 cells indicating the positively and negatively selected guides following the selection with MRTX849 (500 nm) and RMC-6236 (7.5 nM). (C) Impact of MRTX849 and RMC-6236 on the normalized counts of the individual guides targeting CDK2 in UM53 cells based on CRISPR-CAS9 screening. (D) Effect of CDK2 depletion using siRNA on the proliferation of UM53 and MiaPCa-2-MR cells in the absence and presence of MRTX849 (500 nM). Error bars were calculated from mean and SD from triplicates. *** p<0.0001 as determined by 2-way ANOVA. Experiments were done at 3 independent times. (E) Immunofluorescence assay to examine the CDK2 expression in UM53-WT and CDK2-del cells. Scale bar represents 75 microns. (F) Western blotting for CDK2 expression in UM53-WT and CDK2-del cells. (G) Differential effects of MRTX849 and RMC-6236 on the proliferations of UM53-WT and CDK2-del cells. Error bars represent mean and SD from triplicates. Experiments were repeated three independent times. *** p<0.0001 as determined by 2-way ANOVA. (H) Impact of CDK2 deletion on cellular outgrowth of UM53 cells following removal of RMC-6236 after 96 hours of treatment. Error bars represent mean and SD from triplicates. *** p<0.0001 as determined by 2-way ANOVA.

To investigate whether additional cell cycle regulators modulate the response to KRAS inhibition, CRISPR screen data was analyzed from UM53 cells treated with the KRAS inhibitor MRTX849 [22]. Among all CDKs evaluated, *CDK2* emerged as the most prominent gene whose loss enhanced sensitivity to KRAS inhibition (Fig. 5B). A parallel screen performed in the presence of the pan-RAS inhibitor RMC-6236 revealed a similar pattern, with *CDK2*-targeting guides showing the strongest depletion compared to guides targeting other CDKs (Fig. 5B). Examining the individual guides revealed that three independent *CDK2* sgRNAs were comparably depleted following treatment with MRTX849 and RMC-6236 (Fig. 5C). To validate these findings, *CDK2* was depleted using gene-specific siRNA in resistant UM53 and MiaPaCa-2-MR cells, which significantly enhanced the antiproliferative effects of KRAS inhibition, thereby overcoming resistance (Fig. 5D). As an orthogonal approach, a stable CDK2 knockout line (UM53-*CDK2*-del) was generated using CRISPR-Cas9. Efficient deletion of *CDK2* was confirmed at the protein level by both immunofluorescence and immunoblot analysis, demonstrating robust loss of CDK2 expression in UM53 cells (Fig. 5E, F). *CDK2* deletion significantly potentiated the efficacy of both MRTX849 and RMC-6236 (Fig. 5G). Additionally, *CDK2* deletion significantly delayed the cellular outgrowth in UM53 cells following the removal of RMC-6236, suggesting that CDK2 is required for efficient cell cycle re-entry after RAS inhibitor-induced arrest (Fig. 5H).

### Co-targeting CDK2 and KRAS to overcome resistance

Given these genetic data indicating that loss of CDK2 enhances the efficacy of KRAS inhibition, we next examined whether pharmacologic inhibition of CDK2 catalytic activity would provide similar results. While the recently developed CDK2-selective inhibitors are under clinical investigation in other malignancies including ovarian and breast cancers, we evaluated the therapeutic potential of co-targeting CDK2 and KRAS (CDK2i+KRASi) in KRAS-mutant cancer models [36]. In resistant models, including MiaPaCa-2-MR, UM53 and AsPC-1-MR cells, combined treatment with the CDK2 inhibitor INX-315 and the mutant-selective KRAS or the RAS-ON inhibitor, resulted in cooperative suppression of cell proliferation compared to single-agent treatments (Fig. 6A). The combination of INX-315 and MRTX849 also resulted in an enhanced inhibition of spheroid growth, derived from MiaPaCa-2-MR cells (Fig. 6B). These findings indicate that pharmacologic CDK2 inhibition restores sensitivity to both mutant-specific and pan-RAS inhibition.

**Figure 6:**
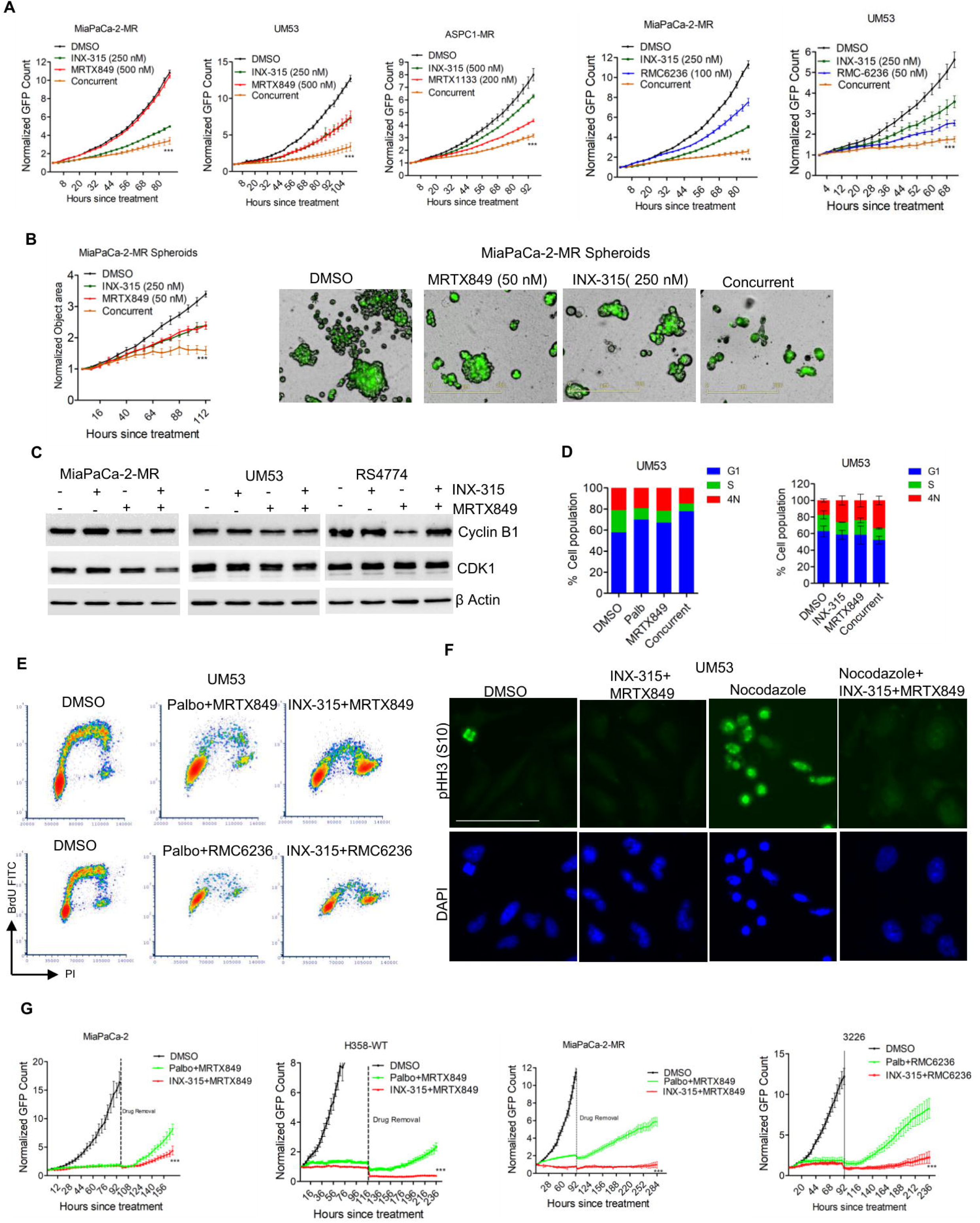
Cellular outcomes of co-targeting CDK2 and KRAS/RAS: (A) Live cell imaging to monitor the proliferation of MiaPaCa-2-MR, AsPC-1-MR and UM53 cells following the treatment with INX-315 in combination with mutant-specific KRAS inhibitors. The effect of INX-315 in combination with RMC-6236 on the proliferation of MiaPaCa-2-MR and UM53 cells. Error bars represent mean and SD from triplicates. *** represents p<0.0001 as determined by 2-way ANOVA. (B) The impact of INX-315 in combination with MRTX849 on the growth of spheroids derived from MiaPaCa-2-MR cells. Error bars were determined from mean and SEM from triplicates. *** represents p<0.0001 as determined by 2-way ANOVA. Experiment was done at 3 independent times. Representative images of the spheroids from MiaPaCa-2-MR cells. (C) Western blotting on the indicated proteins from MiaPaCa-2-MR, UM53 and RS4774 cells following the treatment with INX-315 in combination with MRTX849 up to 48 hours. (D) Stack plots illustrating the % of cell population at each phase of cell cycle based on PI profile following the treatment with MRTX849 in combination palbociclib or INX-315. (E) Bivariate flow cytometry analysis of UM53 cells showing the effects of palbociclib or INX-315 in combination with MRTX849 or RMC-6236 on BrdU incorporation. (F) Immunofluorescence analysis of phospho–histone H3 (S10) in UM53 cells pretreated with DMSO or INX-315 in combination with MRTX849 for 48 hours and subsequently treated with nocodazole (250 nM) for 24 hours. Scale bar represents 75 microns. (G) Comparison of cellular outgrowth in the indicated cell lines following removal of KRAS or RAS inhibitors in combination with palbociclib or INX-315 after 4 days of treatment. Error bars represent mean and SD from triplicates. *** represent p<0.0001 as determined by 2-way ANOVA.

To assess the mechanistic impact on cell cycle regulation, the effects of combined CDK2 and KRAS inhibition on key cell cycle proteins were examined. Unlike the combinations involving CDK4/6 inhibition, CDK2 inhibition did not suppress the cell cycle proteins cyclin B1 and CDK1, despite effective suppression of cell proliferation (Fig. 6C). Given that the repression of E2F-target genes are hallmarks of G1 arrest, these findings suggest that CDK2i+KRASi-mediated growth suppression occurs independently of the G1 checkpoint. Consistent with these biochemical differences, cell cycle phase analysis revealed that combined CDK4/6 and KRAS inhibition induced a pronounced G1 arrest, whereas CDK2 and KRAS co-inhibition did not result in accumulation of cells at a discrete cell cycle phase (Fig. 6D). Bivariate flow cytometric analysis demonstrated that combination treatment of palbociclib with MRTX849 or RMC-6236 resulted in accumulation of cells in the G1 phase, thereby preventing entry into S phase, as evidenced by reduced BrdU incorporation (Fig. 6E). In contrast, INX-315 in combination with KRAS/RAS inhibitors resulted in cells entering S phase, with limited BrdU incorporation, indicating impaired DNA synthesis (Fig. 6E). Moreover, the CDK2i+KRASi or RASi increased the population of cells with 4N DNA content indicating an arrest at G2/M phase (Fig. 6E). To more precisely define the fate of cells with 4N DNA content induced by combined INX-315 and MRTX849 treatment, cells were subsequently exposed to nocodazole, a microtubule-disrupting agent that induces mitotic arrest [37]. As expected, nocodazole treatment alone increased phosphorylation of histone H3 (pHH3), a marker for mitotic arrest (Fig. 6F). However, pretreatment with INX-315 and MRTX849 inhibited the nocodazole-induced pHH3 accumulation, indicating that the combination treatment arrests cells at the G2 phase and prevents entry into mitosis (Fig. 6F).

Apart from inhibiting multiple cell cycle processes, the CDK2i+KRASi combination induced a more durable response as compared to the CDK4/6i+KRASi treatment. In multiple models tested (MiaPaCa-2, H358, MiaPaCa-2-MR and 3226) both CDK4/6i+KRASi and CDK2i+KRASi inhibited cell proliferation as an acute response; however, the CDK4/6i combination treatment displayed recovery of cell proliferation four days after treatment cessation, indicating cell cycle re-entry, whereas CDK2i+KRASi treatment significantly delayed cellular outgrowth following drug removal, exhibiting a more durable response (Fig. 6G). Notably, single-agent treatments were substantially less effective in MiaPaCa-2 and 3226 models, as cells resumed rapid proliferation upon drug withdrawal (Fig. S7A). Overall, targeting the cell cycle by inhibiting CDK4/6 or CDK2 enhanced the antiproliferative effects of KRAS and/or RAS inhibitors, resulting in distinct cellular outcomes.

### *In vivo* efficacy of co-targeting KRAS with CDK4/6 and CDK2

To determine whether the *in vitro* findings translate *in vivo*, we evaluated the anti-tumor efficacy of KRAS inhibition in combination with CDK4/6 inhibitor. Xenografts derived from MiaPaCa-2-MR cells and RS4774 patient-derived tumors were established in immunodeficient mice and treated with the KRAS inhibitor MRTX849 and the CDK4/6 inhibitor palbociclib, either as single agents or in combination. While single-agent treatments yielded modest impact on tumor growth, the combination significantly suppressed tumor progression and resulted in more durable disease control in both MiaPaCa-2-MR and RS4774 xenografts (Fig. 7A & 7B). No overt toxicity was observed due to the combination treatment, as no significant change in body weight was observed (Fig. 7C). To assess the *in vivo* mechanistic effects, RB phosphorylation was evaluated by immunohistochemistry in tumor tissues collected from MiaPaCa-2-MR xenografts and RS4774 PDX tumors after 19 and 15 days of treatment, respectively. Single-agent treatment with either palbociclib or MRTX849 resulted in limited effects on RB phosphorylation in both the models (Fig. S8A). However, the combination treatment potently inhibited RB phosphorylation in both MiaPaCa-2-MR xenografts and RS4774 PDX (Fig. 7D). Further biochemical analysis from MiaPaCa-2-MR tumor tissues confirmed the dephosphorylation of RB and suppression of the E2F target cell cycle proteins, cyclin A and cyclin B1 following the combination treatment indicating a G1 cell cycle arrest (Fig. S8B)

**Figure 7:**
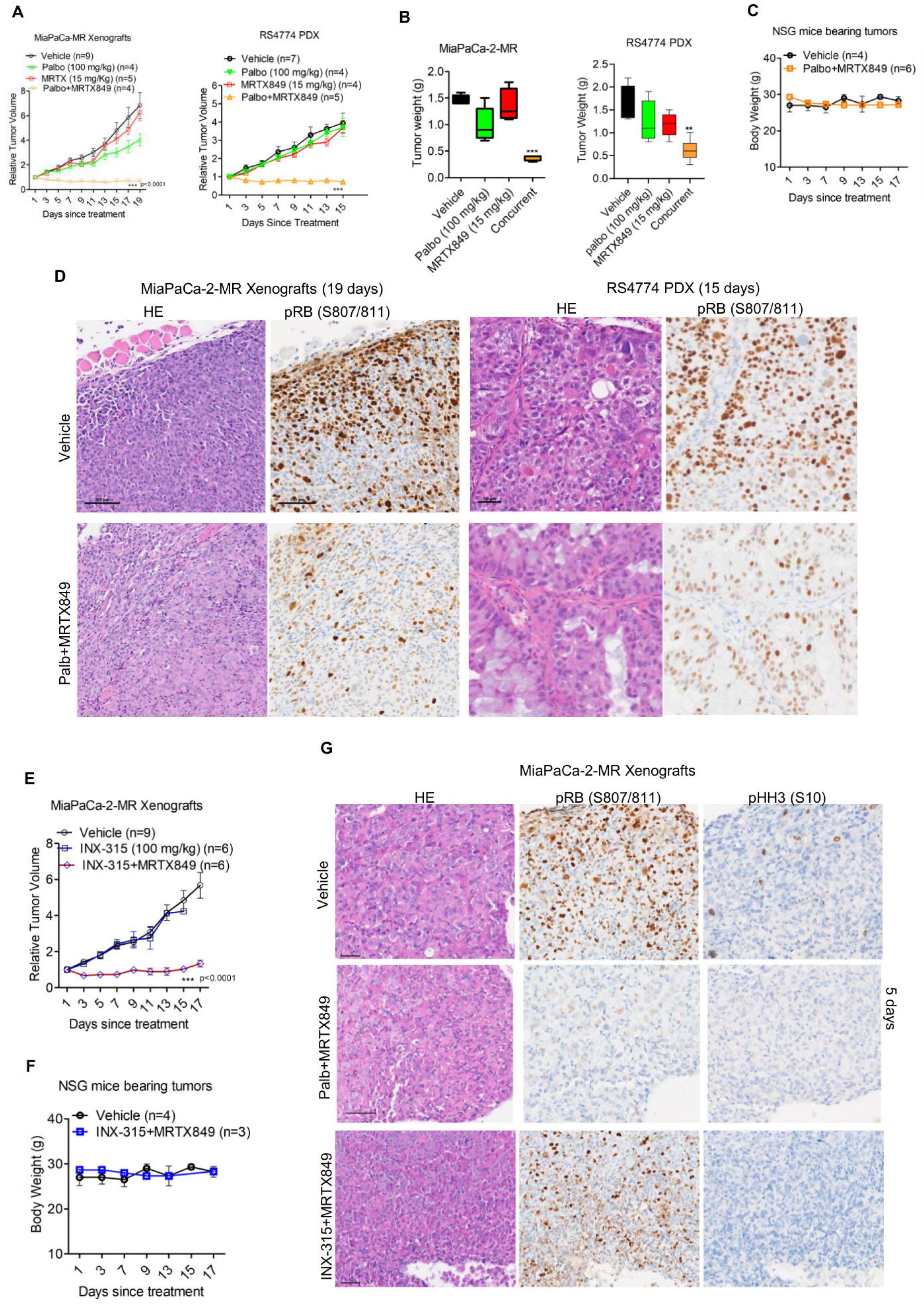
Co-targeting KRAS with CDK4/6 or CDK2 elicit anti-tumor effects: (A) *In vivo* effect of palbociclib in combination with MRTX849 on tumor growth in MiaPaCa-2-MR xenografts and RS4774 PDX. Error bars were determined based on mean and SEM. *** represents p value <0.0001 as determined by 2-way ANOVA. (B) Column graphs illustrate the tumor weights from MiaPaCa-2-MR xenografts and RS4774 PDX following the treatment with palbociclib in combination with MRTX849. *** p<0.0001, ** p<0.01 as determined by unpaired student t-test. (C) Effect of palbociclib in combination with MRTX849 on the mice body weights. Error bar represents mean and SEM. (D) Immunohistochemical analysis of RB phosphorylation in tumor tissues from MiaPaCa-2-MR xenografts and RS4774 PDX models following treatment with palbociclib in combination with MRTX849. Corresponding H&E-stained sections are shown. Scale bar represents 100 microns. (E) Effect of INX-315 in combination MRTX849 on tumor progression in mice bearing MiaPaCa-2-MR xenografts. The change in mice body weights following the combination treatment was monitored. (F) Immunohistochemical analysis of phospho-RB and phospho–histone H3 in tumor tissues following 5 days of treatment with palbociclib or INX-315 in combination with MRTX849. Scale bar represents 100 microns.

We next evaluated whether CDK2 inhibition using INX-315 exerts anti-tumor effects *in vivo* in MiaPaCa-2-MR xenografts. Consistent with the *in vitro* and spheroid findings, combined treatment with MRTX849 and INX-315 significantly suppressed tumor growth without overt toxicity, as indicated by stable body weights (Fig. 7E & 7F). To compare the *in vivo* molecular outcomes of CDK2 and CDK4/6 co-targeting, tumor tissues were analyzed following 5 days of treatment. As observed with prolonged treatment, palbociclib in combination with MRTX849 induced robust suppression of RB phosphorylation, accompanied by reduced histone H3 phosphorylation, consistent with G1 cell cycle arrest (Fig. 7G). In contrast, tumors treated with INX-315 and MRTX849 retained RB phosphorylation despite marked tumor growth inhibition, indicating that the anti-tumor effect was not mediated through G1 arrest (Fig. 7G). Instead, the combination inhibited histone H3 phosphorylation, suggesting impaired mitotic entry (Fig. 7G). These findings are consistent with the cell culture data demonstrating that CDK2 co-targeting disrupts cell cycle progression beyond G1-phase. Collectively, co-targeting KRAS with either CDK4/6 or CDK2 produced strong anti-tumor responses *in vivo*, but through mechanistically distinct effects on cell cycle progression.

## Discussion

The development of mutant-selective KRAS and pan-RAS-ON inhibitors represents a major therapeutic advance, yet accumulating clinical evidence indicates that resistance remains an inevitable challenge [15]. A deeper understanding of the biological mechanisms that enable tumor cells to escape sustained KRAS/RAS pathway suppression is therefore essential to improve the durability of RAS-targeted therapies. Consistent with the clinical emergence of acquired resistance, our study demonstrates that KRAS-mutant PDAC and NSCLC cell lines developed resistance following prolonged exposure to mutant-selective KRAS inhibitors MRTX849 and MRTX1133 *in vitro*, and this phenotype was retained *in vivo* in xenograft models [38–40]. In contrast to patients, where resistance frequently arises through secondary KRAS mutations that impair binding of mutant-specific inhibitors and restore downstream signaling, our resistant models maintained target engagement, as evidenced by sustained suppression of ERK phosphorylation both *in vitro* and *in vivo* [15, 41, 42].

Reactivation of receptor tyrosine kinase (RTK) signaling, including EGFR, FGFR, MET, and RET has also been observed as a bypass mechanism to KRAS inhibition in patients [38, 39]. Notably, transcriptional profiling of patients who progressed on sotorasib (AMG510) demonstrated enrichment of gene programs associated with parallel mitogenic pathways and cell cycle progression, which is consistent with our observations in MiaPaCa-2-MR cells [38]. These findings have provided a rationale for vertical co-targeting strategies combining KRAS and RTK inhibitors [43]. Combining KRAS inhibitors with EGFR-directed therapies has particularly emerged as a potential therapeutic approach in clinical settings. However, given the diversity of compensatory signaling pathways engaged during resistance, targeting any single upstream pathway may be insufficient to restore therapeutic sensitivity in different tumor models [44]. This phenomenon was observed in our systematic combinatorial drug screen across resistant models, which revealed substantial heterogeneity in pathway dependencies, with the MiaPaCa-2-MR cell line failing to exhibit durable response to any upstream co-targeting approach. Given that RTK-mediated signaling is transduced through RAS, we evaluated the pan-RAS inhibitor RMC-6236 as a strategy to broadly suppress compensatory signaling [45]. Although RMC-6236 produced enhanced inhibition of mitogenic signaling and robust antiproliferative effects across resistant models, acquired resistance again emerged following prolonged treatment.

To overcome the limited and heterogeneous responses associated with co-targeting parallel mitogenic pathways, we shifted our focus downstream of KRAS to identify convergent effectors capable of uniformly suppressing proliferation. Transcriptomic and global proteomic analyses revealed that cell cycle programs were differentially impacted by KRAS inhibitors between the sensitive and resistant models. Notably, in resistant cells, cell cycle progression remained active despite effective KRAS/RAS pathway suppression, indicating functional decoupling of proliferative control from upstream mitogenic inputs. Mechanistically, RAS signaling regulates cell cycle progression through modulating the catalytic activities of key CDKs. Oncogenic RAS promotes cyclin D1 expression to activate CDK4/6 and facilitates degradation of the CDK2 inhibitor p27^Kip1^, thereby enabling CDK2 activation [46–48]. Therefore CDK4/6 and CDK2 kinases are considered critical downstream mediators of KRAS-driven proliferation and potential nodes of convergence for therapeutic intervention. Based on this rationale, selective inhibitors of CDK4/6 and CDK2 were used as two independent strategies to suppress proliferative escape and overcome resistance to KRAS/RAS inhibition across models.

Co-targeting CDK4/6 with KRAS inhibition is a well-established therapeutic strategy and is currently being explored in multiple preclinical settings. This approach is supported by prior studies demonstrating that CDK4/6 inhibitors cooperate with agents targeting KRAS-mediated signaling pathways, including MEK, ERK and mTOR, across several cancer types [26, 49–51]. Our data also demonstrated that the CDK4/6 inhibitor palbociclib enhanced the efficacy of KRAS inhibitors, overcame acquired resistance across models, and produced durable disease control *in vivo*. Unlike CDK4/6, CDK2 represents a relatively underexplored therapeutic target in the context of RAS-driven tumors [52]. Mechanistically, combined KRAS and CDK4/6 inhibition converges on inactivating CDK2, positioning it as a critical effector of cell cycle progression. Moreover, our genome-wide CRISPR screen independently identified CDK2 as the most prominent CDK whose loss markedly enhanced sensitivity to KRAS inhibition. Although CDK2 inhibitors have thus far been evaluated in limited clinical settings, our findings establish their therapeutic potential in KRAS-mutant tumors and highlight CDK2 as a promising strategy to overcome RAS inhibitor resistance.

Although both CDK4/6 and CDK2 inhibitors enhanced the antitumor efficacy of KRAS/RAS inhibitors, their cellular consequences were mechanistically distinct. In a canonical cell cycle, CDK4/6 and CDK2 function sequentially to regulate RB phosphorylation and cell cycle progression, leading to the prevailing view that their therapeutic targeting would produce comparable outcomes [53, 54]. However, our data demonstrates that CDK4/6 inhibition primarily enforces a G1 cell cycle arrest, whereas CDK2 inhibition exerts broader control over cell cycle progression, including regulation of DNA replication and mitotic entry [52]. Notably, co-targeting CDK2 with KRAS/RAS inhibition produced more durable suppression of proliferation in selected tumor contexts, although the determinants of this context-specific sensitivity remain to be defined. In conclusion our study indicates that co-targeting CDK4/6 and KRAS or RAS produces a robust and more uniform anti-proliferative response across resistant models. In contrast to strategies aimed at upstream mitogenic signaling, which exhibit substantial context dependence, cell cycle targeting represents a convergent and broadly effective approach to overcoming resistance to KRAS-directed therapies.

## Acknowledgements

The authors thank all members of the laboratory group and colleagues in the discussion and preparation of the manuscript. We thank Incyclix Bio for providing INX-315. This work was supported by grants from the National Institutes of Health, National Cancer Institute (CA267467, CA211878 and R37CA275961), Roswell Park Alliance Foundation (GR75001546) and S10 Instrumentation Award (S10OD030410). Moreover, this work was supported by the National Cancer Institute (NCI) grant P30CA016056 involving the use of Roswell Park Comprehensive Cancer Center’s Advanced Tissue imaging (ATISR), Drug Discovery Core (DDCSR), Experimental tumor models (ETM) and Genomics shared resources. The content is solely the responsibility of the authors and does not necessarily represent the official view of the National Institutes of Health. The article was proofread by Thomas O’Connor.

## Author contributions

Study concept and design: VK, EA, ESK and AKW

Acquisition of data: VK, JW and EA

Analysis and interpretation of data: VK, JW, EA, ESK and AKW

Study supervision: VK, ESK and AKW.

