## Supplementary information for "Targeting Distinct Cell Cycle Nodes Overcomes KRAS/RAS Inhibitor Resistance"

Fig S1

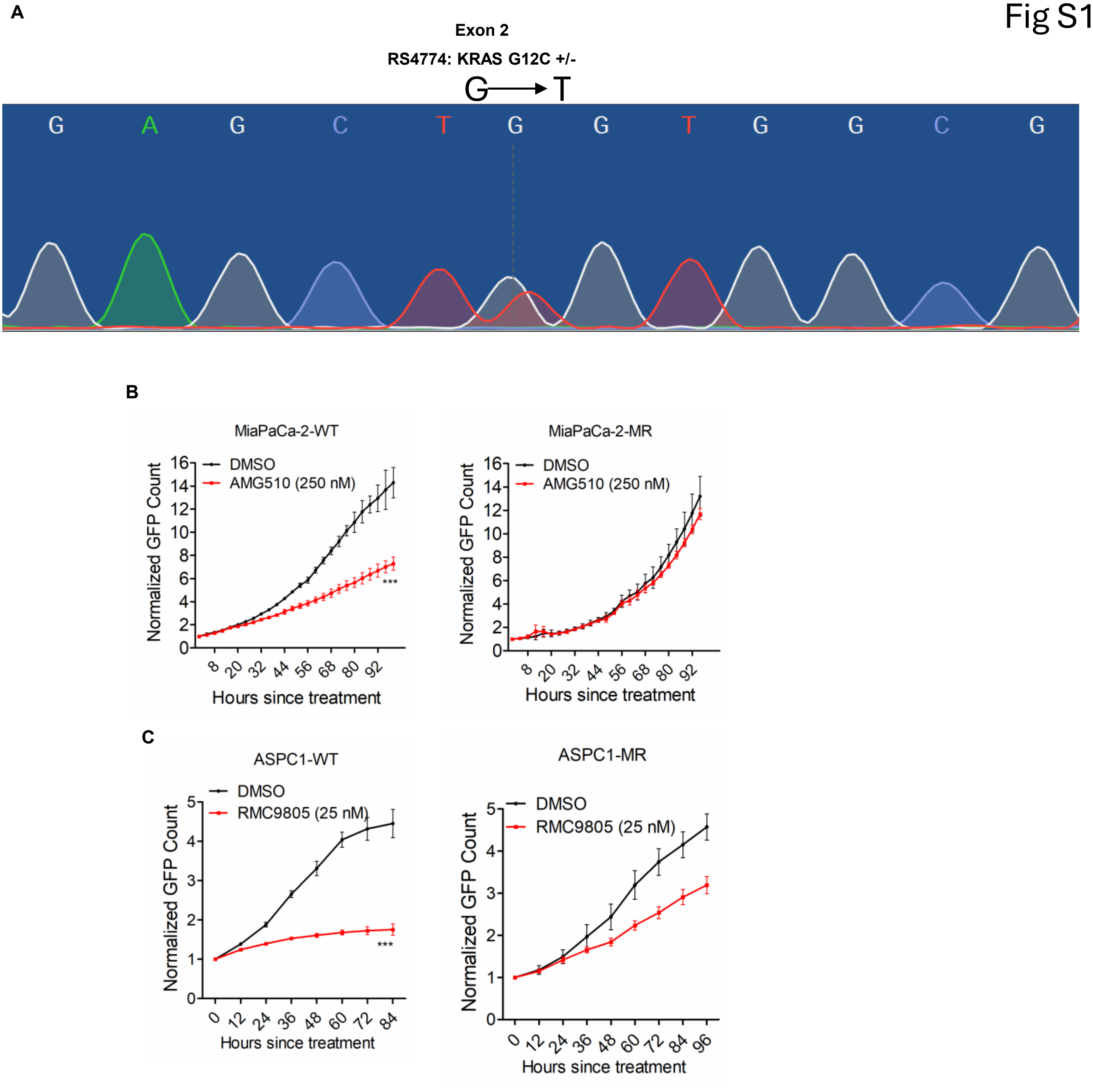


(A) Sanger sequencing of PCR-amplified KRAS exon 2 from genomic DNA of RS4774 cells, demonstrating a heterozygous G>T substitution, as indicated by overlapping peaks at the mutation site. (B) Differential effect of AMG510 on the proliferation of MiaPaCa-2-WT and MiaPaCa-2-MR cells. Error bars represent mean and SD from triplicates. Experiment was done at 3 independent times. *** represents $p<0.0001$ as determined by 2-way ANOVA. (C) Live cell imaging on ASPC1-WT and ASPC1-MR cells following the treatment with mutant-specific KRAS inhibitor, RMC-9805. Error bars represent mean and SD from triplicates. Experiment was done at 3 independent times. *** represents $p<0.0001$ as determined by 2-way ANOVA.

### Fig S2

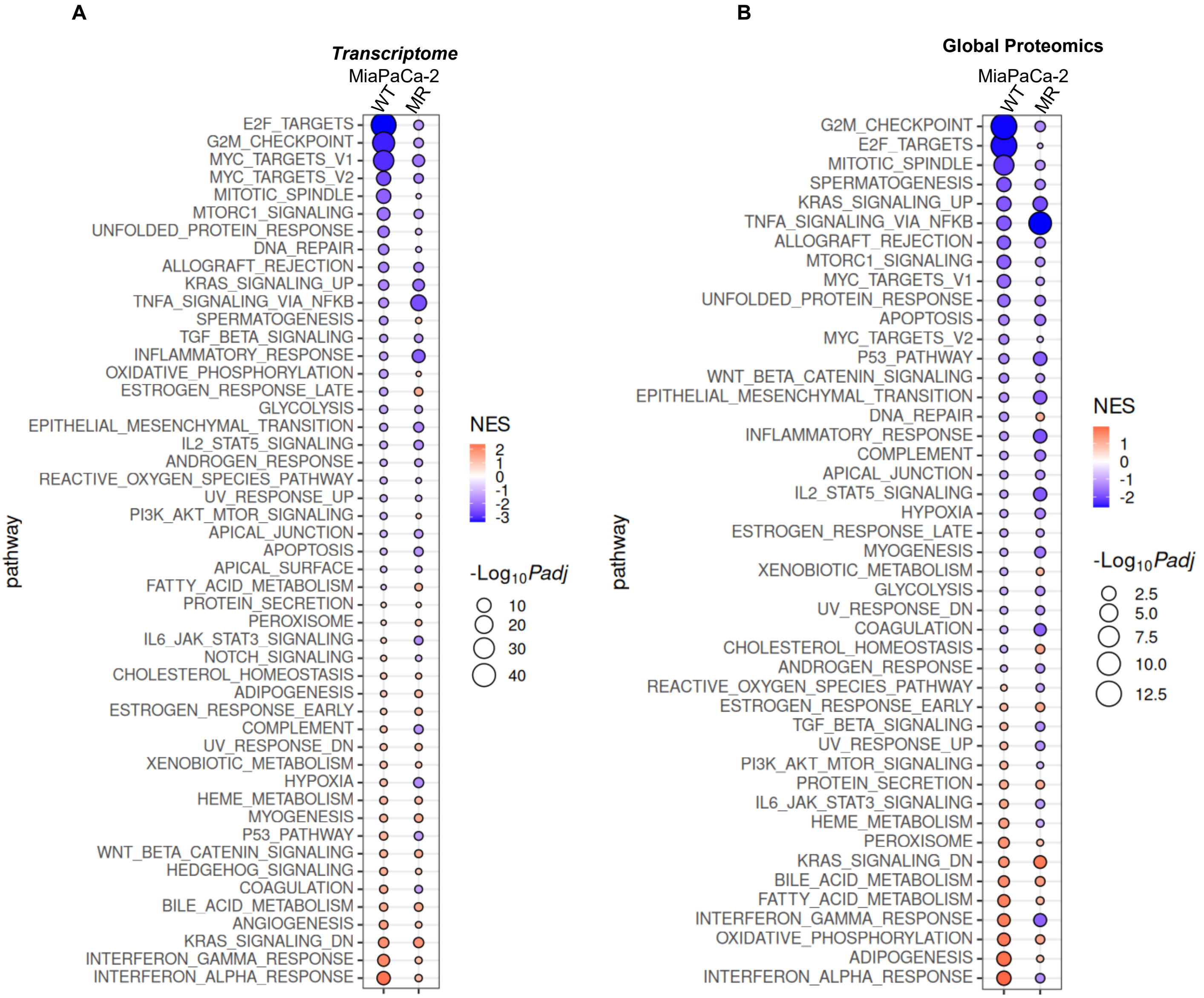


(A) Bubble plot showing differentially enriched pathways in MiaPaCa-2-WT and MiaPaCa-2-MR cells following treatment with MRTX849, ranked by gene set enrichment score and statistical significance (P value). **(B)** Gene set enrichment analysis (GSEA) of differentially expressed proteins showing associated pathways ranked by gene set enrichment score and P value.

Fig S3

**A**

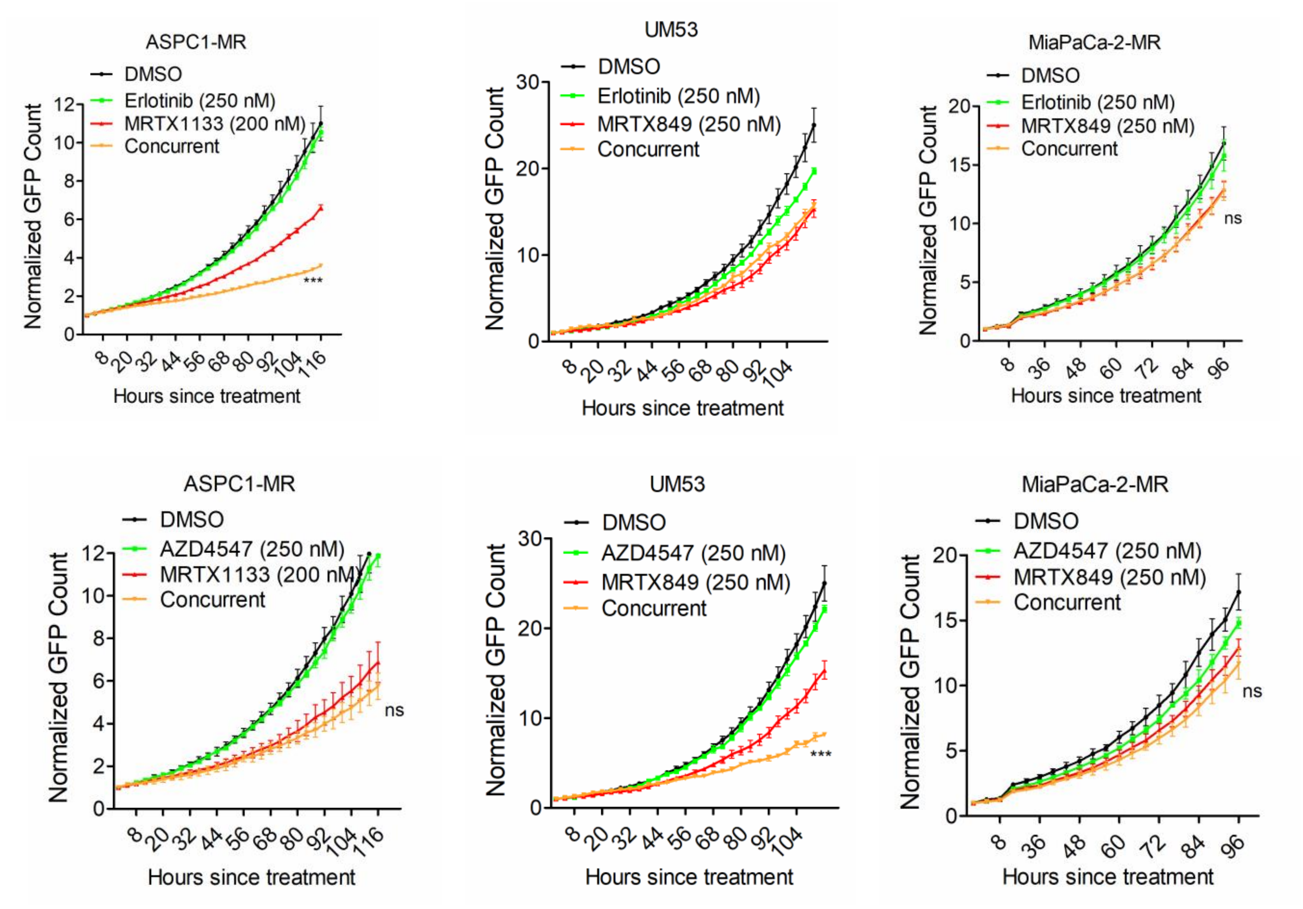


(A) Effect of MRTX1133 in combination with erlotinib or AZD4547 on proliferation of ASPC1-MR, UM53, and MiaPaCa-2-MR cells. Error bars represent mean and SD from triplicates. Experiment was done at 3 independent times. *** represents p<0.0001 as determined by 2-way ANOVA.

Fig S4

A

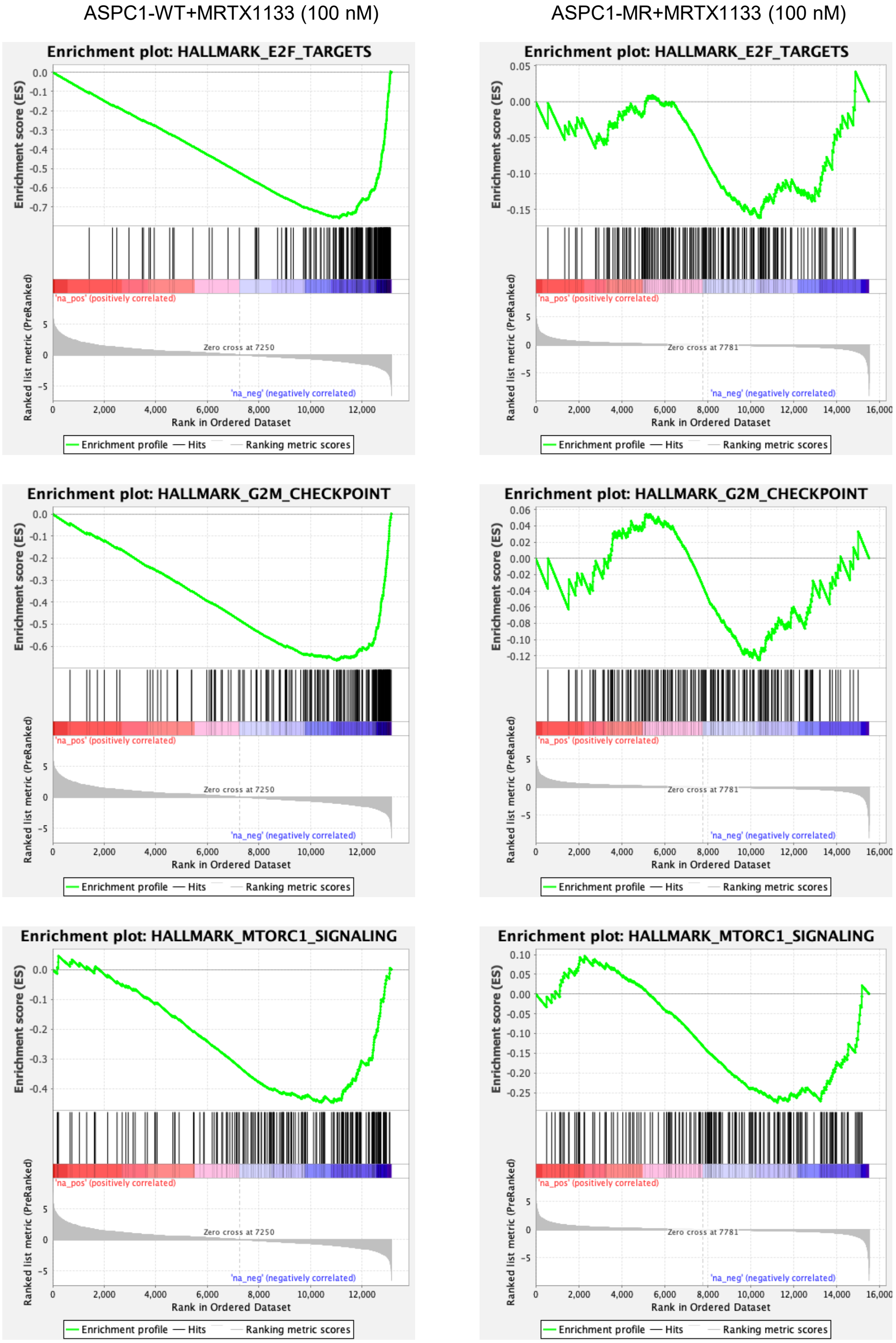


(A) GSEA analysis to demonstrate the differential effect of MRTX1133 and RMC-6236 on the indicated pathways in ASPC1-MR cells.

Fig S5

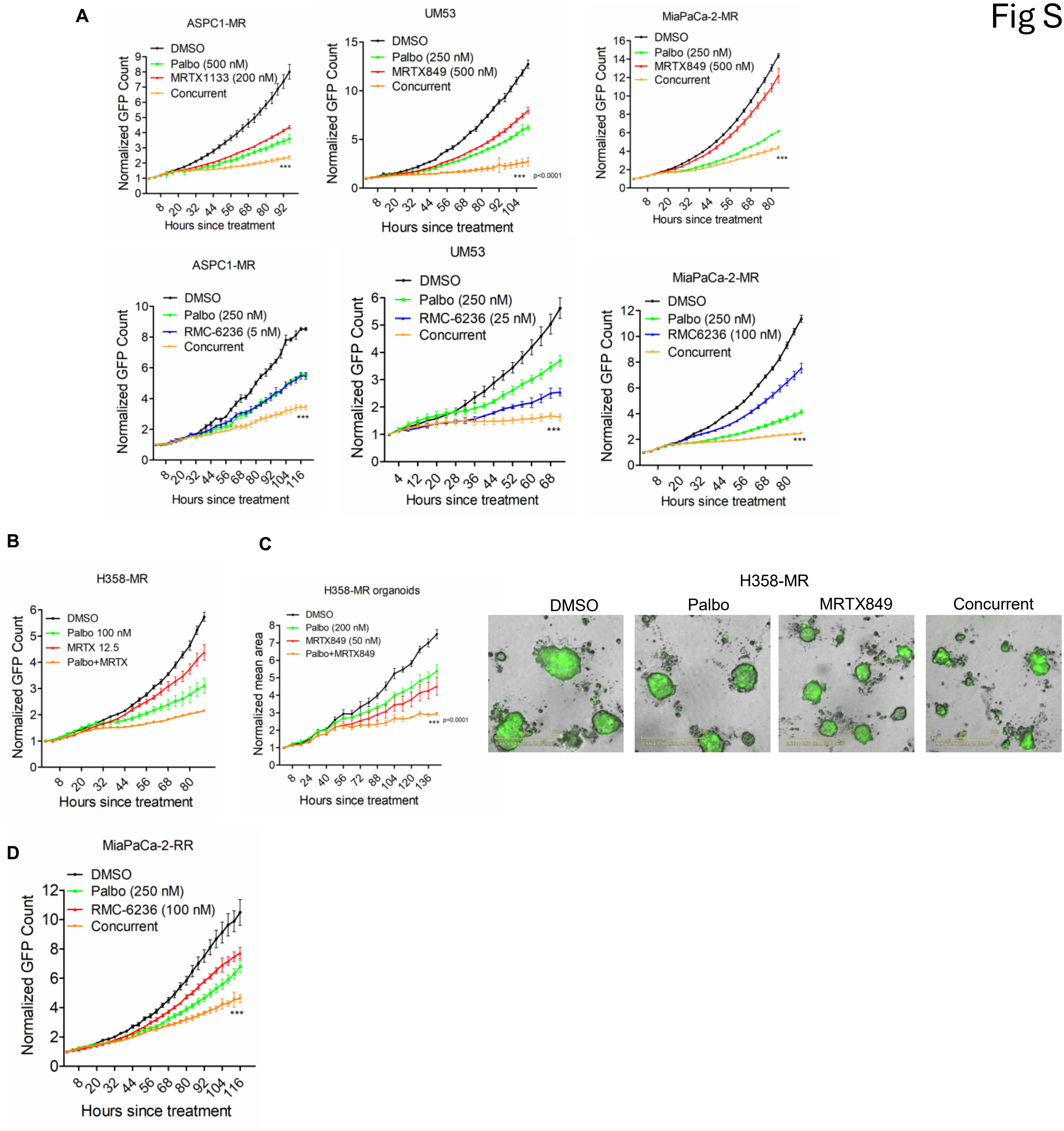


(A) Effect of palbociclib in combination with MRTX1133 or RMC-6236 on proliferation of ASPC1-MR, UM53, and MiaPaCa-2-MR cells. Error bars represent mean and SD from triplicates. Experiment was done at 3 independent times. *** represents p<0.0001 as determined by 2-way ANOVA. (B) Effect of palbociclib in combination with MRTX849 on the proliferation of H358-MR cells. Error bars were calculated from triplicates. *** represents p<0.0001 as determined by 2-way ANOVA. (C) Effect of palbociclib in combination with MRTX849 on the growth of spheroids derived from H358-MR cells. Error bars were calculated from triplicates. *** represents p<0.0001 as determined by 2-way ANOVA. (D) Effect of palbociclib in combination with RMC-6236 on the proliferation of MiaPaCa-2-RR cells. Error bars represent mean and SD from triplicates. Experiment was done at 2 independent times.

### Fig S6

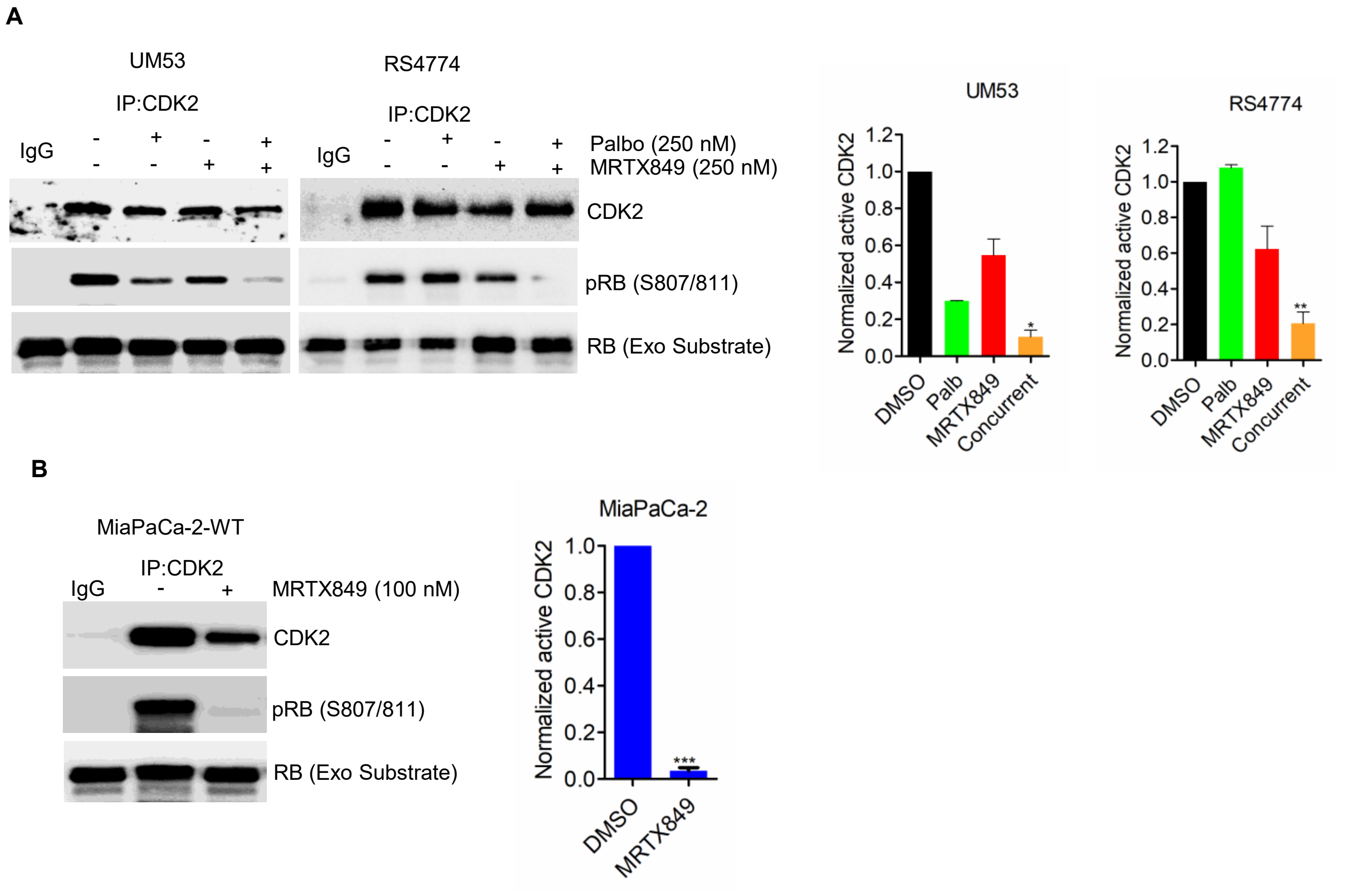


(A) *In vitro* kinase reaction in UM53 and RS4774 cells treated with Palbociclib in combination with MRTX849. CDK2 was immunoprecipitated and subjected to kinase reaction. The phosphorylation of the C-terminal RB peptide fragment was determined by western blotting. Densitometric analysis of RB phosphorylation corresponds to the CDK2 kinase activity. Error bars represent mean and SEM from 3 independent experiments. (B) *In vitro* kinase reaction in MiaPaCa-2-WT cells following the treatment with MRTX849 (100 nM). Column graphs represent the densitometric quantification of pRB. Error bars were calculated based on mean and SEM from triplicates. *** represents $p<0.0001$ as determined by paired –t-test.

### Fig S7

**A**

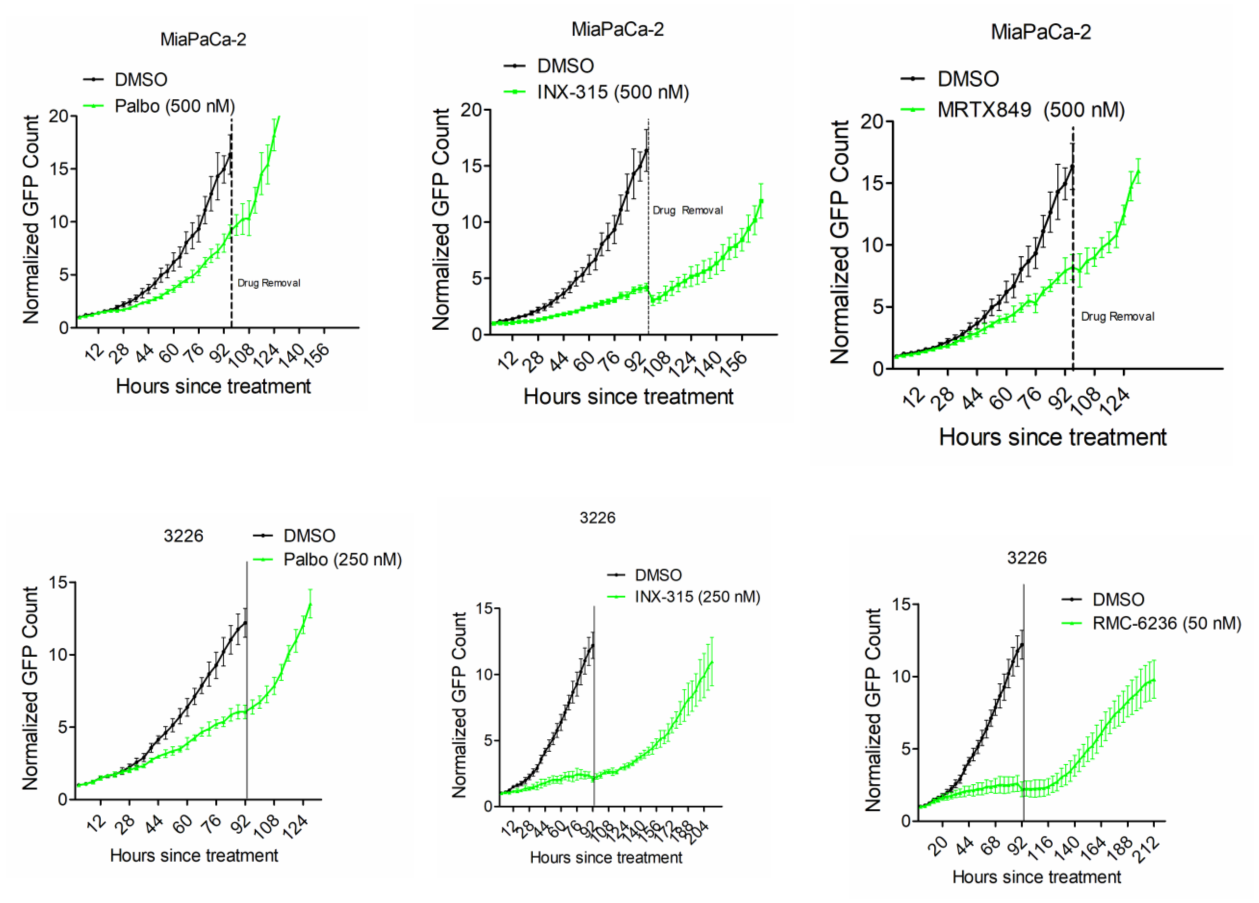


(A) Live cell imaging to monitor the cellular outgrowth in MiaPaCa-2 cells following the removal of palbociclib, INX-315 and MRTX849 after 4 days of treatment. 3226 cells were treated with palbociclib, INX-315 and RMC-6236 for 4 days followed by drug removal. Error bars represent mean and SD from triplicates. Experiment was done at 3 independent times.

Fig S8

A

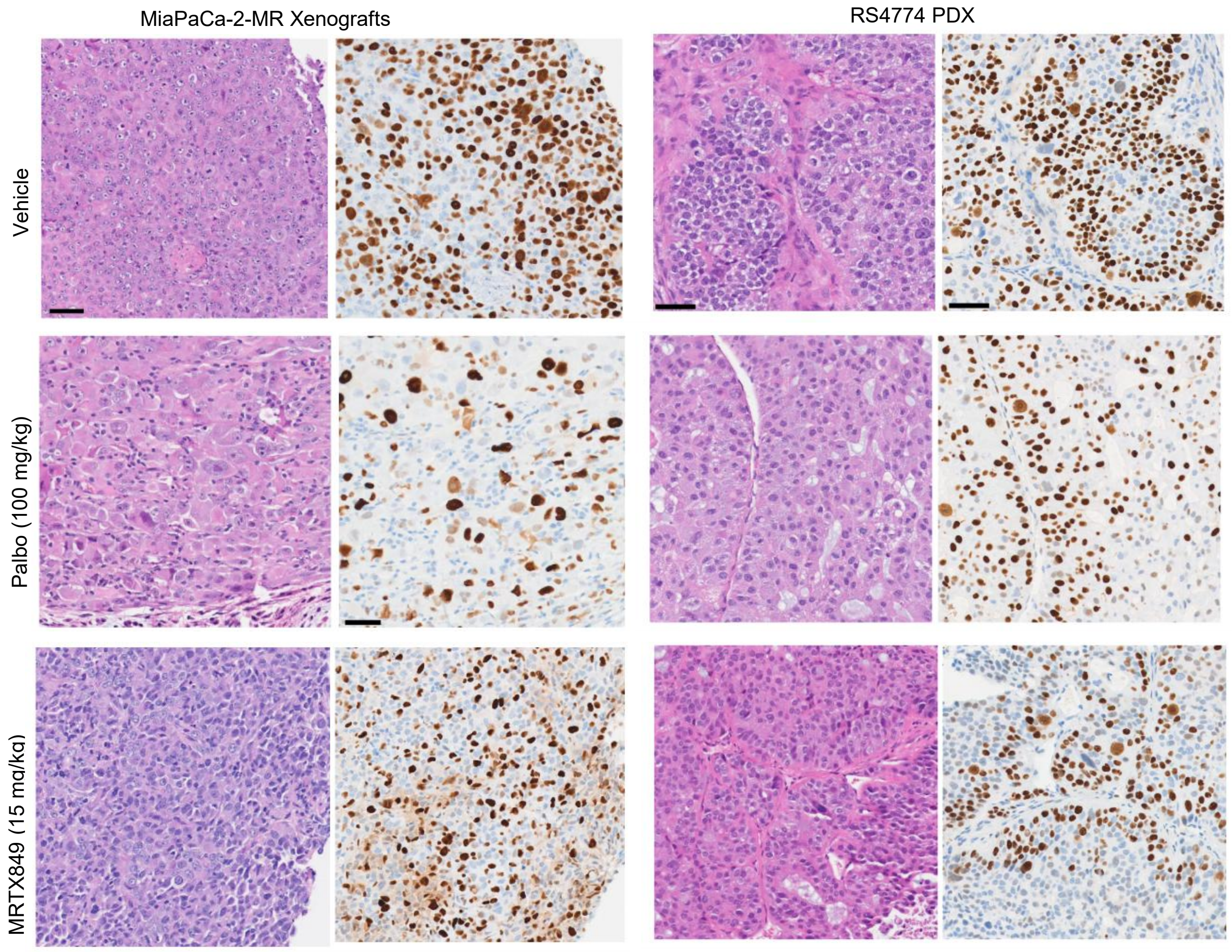


B

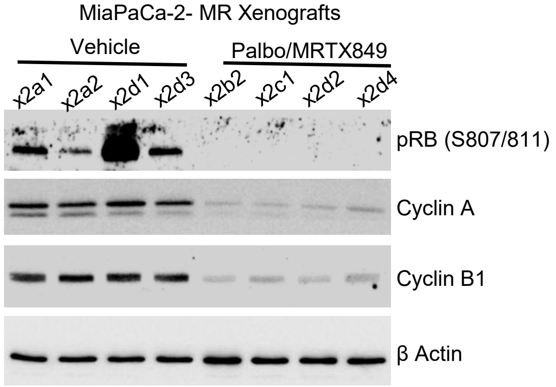


(A) Immunohistochemical staining on tumor tissues excised from MiaPaCa-2-MR xenografts and RS4774 PDX following the treatment with vehicle, palbociclib and MRTX849 to determine the phosphorylation status of RB. The H&E from the corresponding tissues were shown. (B) Biochemical analysis to examine the effect of palbociclib in combination with MRTX849 on the indicated proteins on the tumor tissues excised from MiaPaCa-2-MR xenografts.
